# Active Neural Representation of Stimulus Categories Outside of the Focus of Attention

**DOI:** 10.64898/2026.06.22.733762

**Authors:** Jeffrey D. Johnson, Shawn E. Christ, Nelson Cowan

## Abstract

Previous research on the brain correlates of working memory using functional magnetic resonance imaging (fMRI) and multivariate pattern analysis (MVPA) have shown that neural activity related to information currently needed to respond on a test is considered to be in the focus of attention (FoA). An ongoing controversy pertains to the neural representation of information in working memory that is not needed for the upcoming test but possibly for a subsequent one, which is considered to reside in an activated portion of long-term memory (aLTM). The key theoretical issue regarding aLTM is whether it corresponds solely to an activity-silent neural state. Here, by using a retrospective cuing task in which two arrays of stimuli from different categories were presented in succession on the same trial, and a pattern classifier trained on the first-presented category during the presentation of the second, we provide evidence that aLTM is associated with an active neural state. Specifically, the aLTM effect was demonstrated to be dissociable in direction from that related to the FoA, there was considerable overlap between brain regions representing information in a stronger form in the FoA and a weaker form in aLTM, and the two states appeared to be differentially subject to flexible cognitive control versus natural decay.

## INTRODUCTION

Working memory, the information temporarily held for use in various types of cognitive tasks, appears to include some privileged information held in the focus of attention (FoA) and additional information beyond the FoA. The latter, termed the activated portion of long-term memory (aLTM), is potentially relevant to upcoming task demands and can also include newly-learned associations (Cowan, 1988, 2019; Cowan et al., 2024). Our theoretical questions here are related to how neural representations might differ for working memory information in the FoA versus outside of it, in aLTM.

The extant neuroimaging literature on the FoA and aLTM is inconsistent, and we aim to resolve the inconsistency. Various studies have claimed that the neural representation of information inside the human FoA involves pathways of active neural firing, whereas information temporarily outside the FoA is represented in an activity-silent manner defined by other neural properties such as synaptic weights (Kamiński & Rutishauser, 2020; Lewis-Peacock et al., 2012; Rose, 2020; Rose et al., 2016; Stokes, 2015). In contrast to this claim, however, Christophel et al. (2018) demonstrated neural activity associated with stimuli that were outside the FoA. This discrepancy in the research literature, regarding the nature of information in and out of the FoA (e.g., Iamshchinina et al., 2021), is addressed here with a procedure combining methodological aspects of two previous works (Christophel et al., 2018; Lewis-Peacock et al., 2012). In particular, we used arrays of different stimulus types on the same trial (numbers, colors, and ring orientations), as Lewis-Peacock et al. did, rather than drawing all stimuli from a single type. Additionally, we delivered the separate arrays successively, as in Christophel et al., as opposed to presenting them concurrently. This successive presentation procedure allowed us to distinguish neural activity specific to the type of stimulus currently in aLTM from that associated with stimuli in the FoA.

In the first of these two forerunners of our study, Lewis Peacock et al. (2012) developed a key method using functional magnetic resonance imaging (fMRI) with multi-voxel pattern analysis (MVPA; see Haxby et al., 2014; Haynes & Rees, 2006; Norman et al., 2006) in which two modalities of stimuli were presented on every trial and a third one was omitted from the trial. In their most directly relevant Experiment 2, the modalities were phonological (pseudowords), semantic (words), and visual (line orientations). Two retrospective cue-and-test cycles occurred per trial, either testing both presented modalities in turn (switch trials) or testing two different aspects of the stimuli presented in only one modality (repeat trials). The results showed that upon receiving a cue indicating which modality would be tested, the MVPA classifier evidence for neural activity associated with that modality increased or stayed elevated, whereas the classifier evidence for the other presented modality returned to a baseline level, equivalent with the unpresented third modality. On switch trials, activity for the modality that was not tested with the first cue was found to return from baseline to an elevated level when the second cue indicated that it was about to be tested. Behavioral performance for that second-cued modality was also well above chance. These results suggested that information needed later in the trial had not vanished when its MVPA signal disappeared but rather, until the second cue, it had gone into an activity-silent state with no discernable neural pattern. Rose et al. (2016) subsequently used transcranial magnetic stimulation (TMS) to show that the information that was not in the FoA could be artificially reactivated when it was potentially still needed on the trial (after the first cue) but not when it was understood to be no longer needed (after the second cue), consistent with the idea that activity-silent information is somehow governed by task goals.

In the second forerunner of our study, Christophel et al. (2018) came to conclusions different from those of the aforementioned studies. They presented gratings of two different visual orientations in succession, one to the left and the other to the right of fixation, and then, like in the previous work, used two cues in succession to allow for switch and repeat trials. During an 8-s retention interval after the first cue, fMRI data acquired during this procedure were analyzed using MVPA. The results specific to the orientation of each stimulus showed different non-null patterns for the cued information (presumably in the FoA) versus the uncued information (presumably not in the FoA). Whereas attended information was discernable in separate analyses for three regions of interest—occipital cortex (OC), intraparietal sulcus (IPS), and the frontal eye fields (FEF)—effects for the unattended information were observed in only the latter two regions. The authors concluded that representations of unattended information are held in a form that is not activity-silent, in contrast to the conclusions of Lewis-Peacock et al. (2012) and Rose et al. (2016), though they might be less sensory-related or of lower resolution than the representations in OC for attended information.

There are limitations of the work by Christophel et al. (2018) that leave it open to multiple interpretations. First, although in previous studies one can easily imagine attending to one category of information and ignoring another (e.g., “line orientations are relevant right now; semantic information is not relevant right now but possibly will be later in the trial”), that may not be the case when the two stimuli are both visual orientations. Gossaries et al. (2018) and Cai et al. (2020) showed that the function of the IPS in such cases is not only to hold multiple stimuli in working memory, but also to help maintain distinction between stimuli that are similar. Accordingly, in the Christophel et al. study, it may be that both orientations on a trial must be held in the FoA, even after the first cue, to allow differentiation between them at the time of test. In that case, the differentiation of cued and uncued stimuli need not reflect attention per se, but rather the different roles that these stimuli play in the trial. Moreover, while the uncued information might be still attended by the IPS and FEF, failure to observe activity in OC related to the uncued condition could be due to that information being suppressed or somehow transformed.

To address the question of whether unattended information is held in an activity-silent form or a non-silent form, we reverted to stimuli more like Lewis-Peacock et al. (2012), using three categories: color patches, numbers (single digits), and rings that were oriented in different directions. On a given trial, arrays of stimuli from two categories were presented, as shown in Figure 1a. Unlike Lewis-Peacock et al. and more like Christophel et al. (2018), though, the two arrays were presented in succession rather than concurrently. Importantly, this allowed us to use an interval upon presentation of the second array to retroactively train an MVPA classifier on the stimuli presented earlier in the first array. Given the capacity limitations of the FoA (Cowan, 2001, 2011, 2019; Cowan et al., 2011), the main assumption underlying this classification was that the stimuli from the first array had to leave the FoA for encoding of the second array to proceed. For the category presented first on a trial, accurate classification during that initial array period was taken to reflect information in the FoA. In contrast, a classifier trained on neural patterns from the second period, based on the category label of stimuli presented during the first array, was thus taken as a reflection of information outside of the FoA but in aLTM. We were then able to follow the fate of these classifiers throughout later phases of the trial, which included two cue-and-test cycles corresponding to switch and repeat trials. Although a straightforward expectation was to replicate Lewis-Peacock et al. for the cued information in the FoA, the questions of primary interest were related to how FoA and aLTM patterns for the same category might be simultaneously held in opposition to one another and whether a neural signature of aLTM for uncued information would re-emerge whenever information left the FoA. Additionally, we were able to test whether any such aLTM signal would fail to return when it reflected information no longer needed on the trial, after the second cue, as would be expected on the basis of Rose et al.’s (2016) findings.

**Figure 1.**
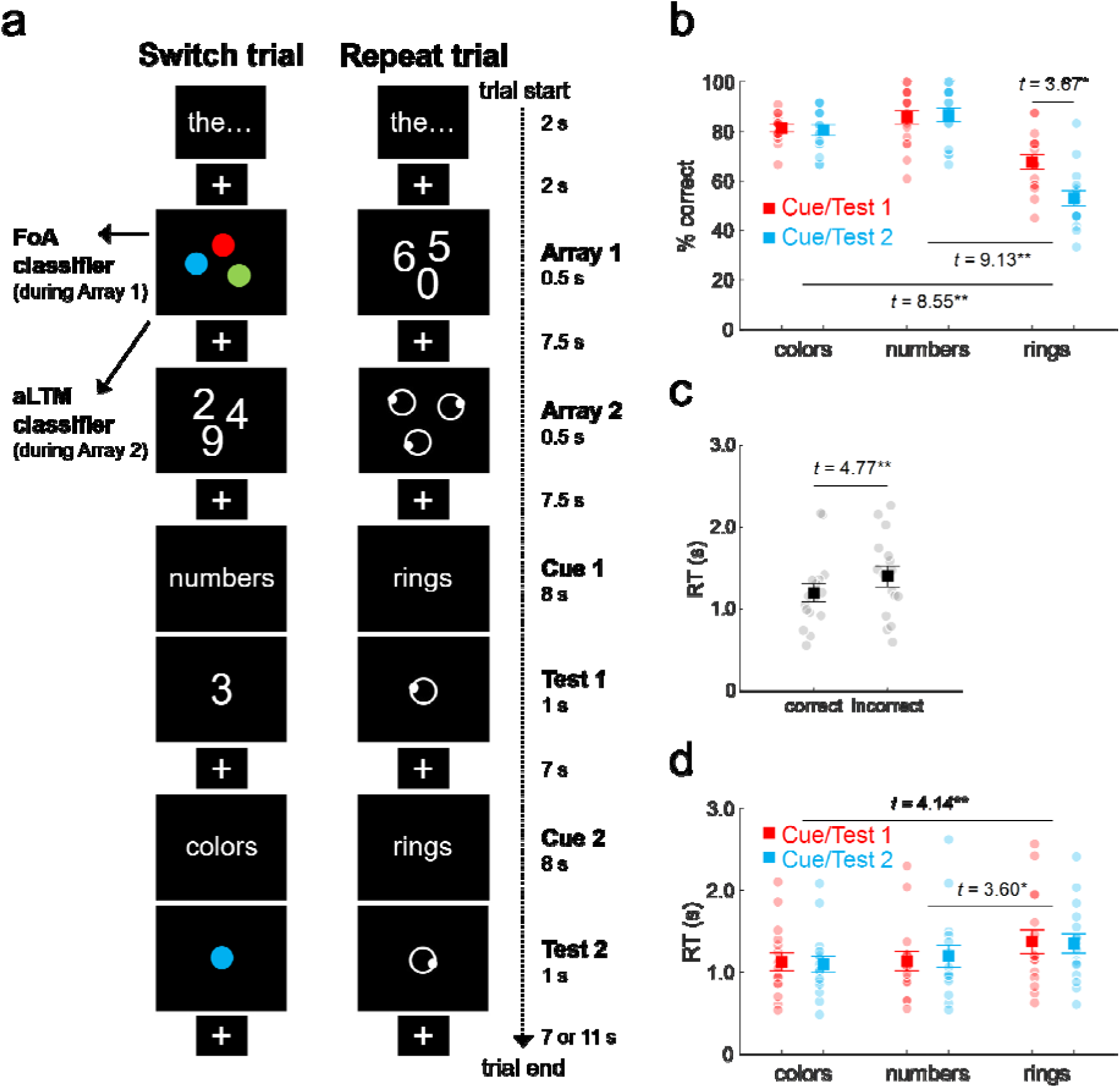
Trial schematic and behavioral results. **(a)** Example stimuli, event sequence, and timing for both the switch and repeat trial types. **(b)** Percentages of correct responses according to stimulus category and whether the category was cued and tested first vs. second. For this and subsequent panels, solid dots and error bars indicate group mean±SEM, and transparent dots correspond to individual subject means. **(c)** Response times (RTs) for correct vs. incorrect recognition judgments. **(d)** RTs for correct judgments, according to stimulus category and whether the category was cued/tested first or second.

## MATERIALS & METHODS

### Subjects

The target sample size of 16 subjects was based on exceeding that of similar studies of working memory that used MVPA with fMRI data (12 subjects in Lewis-Peacock et al., 2015; 10 subjects in Rose et al., 2016). Although our sample size might be considered small compared to other studies, such as Christophel et al. (2018; n = 87), statistical power was maximized by using highly-distinct stimulus categories (cf. the visual gradients of Christophel et al.) and a slow event-related design, along with conducting classification analyses on an individual-subject basis.

Eighteen volunteers from the University of Missouri (MU) community participated in exchange for course credit. All subjects reported being right-handed, learning English as their first language, and having normal or corrected-to-normal vision, no history of neurological disorders, and no other MRI contraindications. Data from two subjects were excluded from all analyses, as one did not complete the experiment and the other responded with chance accuracy (50% correct) on the recognition task. (For comparison, the remaining subjects had an overall correct response rate of 76% [SD = 6; range: 66-88%].) The final sample of 16 (7 females and 9 males) were between the ages of 18 and 22 (M = 19.9) years of age. Informed consent was obtained from all subjects in accordance with the MU Health Sciences Institutional Review Board.

### Stimuli

Examples of the stimuli are provided in the trial schematics in Figure 1a. Stimuli consisted of eight exemplars from each of three categories: colors, numbers, and orientations (rings). Colors were presented as circular, solid patches in the following RGB colors (common descriptors are also provided here): 255,0,0 (red); 0,255,0 (lime green); 0,0,255 (dark blue); 255,255,0 (yellow); 255,0,255 (magenta); 0,255,255 (aqua); 255,127,0 (orange); and 127,127,255 (slate blue).

Numbers were the single digits 0 through 9, excluding 1 and 7 due to visual similarity, and were presented in white 36-point Arial font. Rings consisted of white open circles (filled with black, matching the background) with a smaller white filled circle along the perimeter. The eight ring exemplars were constructed by positioning the smaller circle at 23° (where 0° corresponds to the right of center and moving clockwise) and at intervals spaced every 45°.

At different times during the experiment, stimuli were presented either singularly at the center of the display or as an array of three different exemplars from a single category. For the arrays, stimuli were positioned at eight possible locations around the perimeter of an imaginary circle, starting at 23° and spaced every 45° (0° = right; moving clockwise). At specific times (see below), additional visual stimuli were presented centrally and in white 36-point Arial font, including the category names (‘colors’, ‘numbers’, and ‘rings’), reminders to engage in articulatory suppression (‘the…the…’), and fixation markers (‘+’). All stimuli appeared on the black background of a screen positioned at the head of the magnet bore and viewed through a mirror placed in front of subjects’ eyes. Stimulus presentation and response acquisition were controlled by the Cogent 2000 toolbox in MATLAB (The MathWorks, Natick, MA).

### Design and procedure

Subjects received instructions and completed a short practice version of the experiment prior to entering the MR scanner. Once in the scanner, subjects underwent an anatomical scan (∼8 minutes) and then completed 72 trials of the working memory task. Trials were organized into three blocks of 24 trials. Short breaks were given between blocks and at the midpoint of each block (i.e., after every 12 trials or about every 11 minutes). Scanning was stopped for each break, resulting in six separate fMRI runs. Following completion of the working memory task, the subject was removed from the scanner and debriefed.

See Figure 1a for examples of the stimuli and trial sequence. Two categories of stimuli were used on each trial, resulting in three possible combinations, and the categories could be initially presented in either order. Additionally, as memory was tested twice per trial, each test could be directed at either the category presented first or second, resulting in four test sequences (i.e., first-second, second-first, first-first, and second-second). Blocks of 24 trials contained all possible trial types (3 category combinations × 2 initial presentation orders × 4 testing sequences) presented in random order.

Each trial started with a centrally-presented visual reminder (‘the…the…’ for 2 s) for subjects to engage in articulatory suppression by quietly repeating the word “the” twice per second throughout the trial. A central fixation marker was then displayed for 2 s. An array of three exemplars from a single category appeared for .5 s, followed by a return to the fixation marker for 7.5 s. A second array, consisting of three exemplars from a different category, was then presented with the same timing parameters. A centrally-presented category cue (e.g., ‘colors’) followed and was displayed for 8 s. Subjects were instructed to hold the appropriate stimuli in mind for this entire period. The first recognition test probe then appeared, consisting of a central stimulus displayed for 1 s. Each probe was equally likely to be old or new (randomly chosen in each test instance of each trial). Subjects made the old and new responses with their right index and middle fingers, respectively, by pressing buttons on a response pad held comfortably on their abdomen or lap. A central fixation was displayed for 7 secs before a second category cue, recognition test probe, and fixation proceeded with the same timing. Temporal jitter between trials was introduced by including an additional 4 s of fixation on half of the trials (randomly selected) in each block.

### MRI acquisition and preprocessing

Whole-brain MR data were obtained at the MU Brain Imaging Center on a 3-Tesla Siemens Magnetom TIM Trio scanner equipped with an 8-channel head coil (Siemens Medical Solutions, Erlangen, Germany). T1-weighted anatomical data comprised 176 sagittal slices (256-mm FOV, 1-mm isotropic voxels) acquired with an MP-RAGE pulse sequence. Functional (fMRI) data were acquired using an echo-planar imaging (EPI) pulse sequence sensitive to BOLD contrast (T2*-weighted, 2-sec T_R_, 30-msec T_E_, 90° flip angle). Each fMRI volume consisted of 32 axial slices (3-mm thick, 1-mm gap, ascending interleaved acquisition) with an in-plane resolution of 3×3 mm (192-mm FOV, 64×64 matrix). The fMRI data were acquired in six separate runs (332 volumes each; 12 trials per run).

The data were pre-processed with SPM12 (Wellcome Trust Centre for Neuroimaging, London, UK; http://www.fil.ion.ucl.ac.uk/spm; rev. 6906) in MATLAB (R2020a). After discarding the first four volumes of each run, differences in slice acquisition time were corrected by temporally shifting (via sinc interpolation) the time series of each voxel. Slice-timing correction used the first slice of each TR as the reference, as it corresponded to the onsets of trials and stimuli of interest. The fMRI data were then spatially realigned to the initial volume of the first run followed by a second pass of realignment to the mean volume (across runs). After co-registering the functional and structural data, the structural data were segmented into gray and white matter and deformed to a set of standard tissue probability maps (International Consortium for Brain Mapping; http://www.loni.ucla.edu/ICBM/; Ashburner and Friston, 2005). The data were then normalized with the deformation parameters determined by the previous step, with the fMRI data being resampled into 3-mm isotropic voxels. The data were not spatially smoothed, in contrast with typical pre-processing for univariate fMRI analysis.

### fMRI analysis

After preprocessing, the fMRI data were submitted to a GLM that included six regressors corresponding to the translation and rotation movement parameters and five regressors capturing constant differences across acquisition runs. The GLM also employed a 128-second high-pass filter and an implicit masking threshold of .8 of the global mean activity. Following the modeling, the residual activity for each voxel was z-scored within each run, and the resulting whole-brain maps constituted the patterns used for the classification analyses.

### Whole-brain classification

The fMRI analyses were conducted using the Princeton MVPA toolbox (The Princeton Neuroscience Institute, Princeton, NJ; https://github.com/princetonuniversity/princeton-mvpa-toolbox), with additional functionality implemented in SPM8 and custom MATLAB code. The relevant analysis code and the importance and searchlight maps (see below) are available publicly on the Open Science Framework (OSF) website at https://osf.io/uanqc/.

The analyses were conducted on an individual-subject basis and involved fitting three-way (colors vs. numbers vs. rings) L2-regularized logistic regression models to the patterns of brain activity associated with each stimulus category, resulting in the estimation of voxel-wise weights predicting an output for each category. To make use of the standard “peak” of the delayed and expanded hemodynamic response, patterns corresponding to onsets of 6, 8, and 10 s after stimulus onset (0 s) were assigned to the appropriate category (also see Johnson et al., 2009; McDuff et al., 2009), giving rise to three training patterns from each trial contributing to a given classifier. A ridge penalty (λ) of 100 was selected pre-experimentally, based on previous findings from our and other labs (Hastie et al., 2009; Leiker & Johnson, 2015), and used for all analyses to avoid over-fitting. A three-fold cross-validation procedure was employed by training the model on the data from two blocks (24 trials per block) and testing on the remaining block, with the procedure repeated until each block was tested. The block structure provided equal numbers of trials (and patterns) associated to each stimulus category, with the training data comprising 144 patterns (2 blocks × 24 trials/block × 3 patterns/trial) and the testing data comprising 72 patterns, for each cross-validation iteration. Additionally, feature selection was used to minimize the influence of less informative voxels to the model. Feature selection was implemented by restricting the analysis to the 10,000 voxels (roughly 20% of the total voxels) exhibiting the largest F-values across categories, determined separately for the training data of each cross-validation iteration. To help ensure that our pre-experimental choices of classifier (L2-regularized logistic regression) and classifier parameters (penalty and number of selected features) did not unexpectedly lead to increased overfitting, we also conducted a series of post-hoc analyses using different regularization penalties (both L1 and L2) and feature selection values. The results of these additional analyses are summarized in Figure S1 of the Supplemental Material. Importantly, each of these analyses produced results comparable to those based on our pre-experimental choices.

To separately analyze brain activity associated with the FoA vs. aLTM, we assigned patterns for model training from distinct parts of each trial. FoA-related patterns came from the array of three exemplars that were presented first on the trial (during 0.5 s of presentation and 7.5 s fixation period) and were assigned to the category that was being presented to subjects. By contrast, patterns related to aLTM were taken during the second array of exemplars and following fixation period and were assigned to the category that was presented in the first array because that array was assumed to be removed from the FoA to make room for the newly-encoded second array. For aLTM, this meant that the patterns were assigned to a different category than was currently being presented; importantly, however, the assigned category was equally likely to co-occur with the presentation of exemplars from either of the other two categories. Thus, although there was likely to be stronger activation related to the category presented during the second array, these analyses allowed us to pick up on more subtle activity related to the first-presented category that was in aLTM.

The trained models were evaluated in two ways. First, to determine if the model was successful in identifying category-distinct patterns, they were tested with the data from the left-out block that corresponded to the same time period within the trial. For these analyses, success could be taken as a binary measure of accuracy that was significantly above the chance level (1/3, given the three categories). Second, we evaluated the models according to whether category-related patterns were reactivated during the later parts of the trial (i.e. after two categories had been presented, but subjects were to attend to one of the categories). For these analyses, multiple types of activity (i.e. activity for one category in FoA and for another category in aLTM) were expected to be present at any given time. Thus, accuracy was no longer a useful measure because a single category could not be designated as “correct”. We therefore turned to the model’s raw output, which is a continuous measure for each of the three categories. Importantly, the output (or *evidence*; Lewis-Peacock et al., 2015) could be assessed independently for the two categories present on a given trial and compared to the third, *baseline* category that was not presented on that trial. For both the accuracy and evidence measures, the findings reported here correspond to the average results collapsed across stimulus categories and subjects.

### Searchlight classification

Finally, spatial information about the voxels that were influential to the analyses were assessed by using a “searchlight” selection method (Kriegeskorte et al., 2006; also see, e.g., Johnson et al., 2009). Throughout the entire brain volume, circular searchlights with a radius of two voxels (6 mm) were centered on each voxel, with each resulting volume containing a maximum of 33 voxels (searchlight volumes at the edge of the brain were truncated to exclude non-brain voxels). As with the previous classification analyses, the searchlight analyses also used the three patterns corresponding to the standard peak of the hemodynamic response (6/8/10 s after stimulus onset). Separate L2-regularized logistic regression models (λ = 1, set pre-experimentally) were then run for each searchlight, with the resulting model accuracy assigned to the central voxel. As with the previous analyses for the whole brain (with feature selection), the searchlight analyses were conducted on an individual-subject basis and separately for FoA and aLTM classification.

The searchlight analyses provided the advantage of allowing us to statistically contrast the FoA and aLTM brain maps at the group level. The minimum cluster extent to reach a corrected family-wise error (FWE) rate of *p* < .05 was determined by SPM, individually for the FoA and aLTM searchlight classifiers (vs. chance accuracy). Using a threshold of *p* < .001 (critical *t*_15_ = 3.73), the largest of these extents was 10 voxels (range: 9-10), which was employed for all analyses. Additionally, an inclusive-masking procedure was implemented to identify overlapping results across the FoA and aLTM classifications. Each of the two individual contrasts were thresholded more liberally at *p* < .01, with the resulting conjoint threshold determined to be *p* < .001 according to Fisher’s method of combining probabilities (Fisher, 1950; Lazar et al., 2002). For descriptive purposes, we note that the searchlight analyses gave rise to similar findings as extracting the voxel-wise “importance” values (trained weight × averaged activity; e.g., Johnson et al., 2009; McDuff et al., 2009; Leiker & Johnson, 2015) from the whole-brain classification analyses. All of the results of these analyses, along with the boundaries of the Automated Anatomical Labeling (AAL) atlas (Tzourio-Mazoyer et al., 2002), were mapped onto the smoothed surface of the ICBM152 template brain with BrainNet Viewer (http://www.nitrc.org/projects/bnv/; Xia et al., 2013).

## RESULTS

Examples of the stimuli, along with the sequence and timing of trial events, are displayed in Figure 1a. As an overview, each trial began with successive stimulus arrays (Array 1 and Array 2) from two of three categories (colors, numbers, and rings). Subjects were then cued (Cue 1) to attend to one of the categories, followed by a recognition test (Test 1) on a probe stimulus from that category (equal chance of being presented in the earlier array vs. not). A second cue-and-test sequence (Cue 2 and Test 2) then occurred, either for the same category initially tested (repeat) or for the alternative category (switch). The basic novel finding documented below is as follows: In addition to replicating evidence for information in the FoA, based on a classifier trained on the currently active set during the presentation period, there is independent, novel evidence of activity related to information in aLTM, based on a classifier trained on retention of the first category during presentation of a second.

### Behavioral results

The correct response percentages according to the main manipulations of stimulus category and cue/test order (i.e., first vs. second) are summarized in Figure 1b. (See Figure S2 of the Supplemental Material for the results of additional behavioral analyses involving manipulations that were of lesser importance to our main hypotheses.) An ANOVA of these data gave rise to significant effects of category (*F*_1.9,29.0_ = 53.29, *p* < .001), order (*F*_1,15_ = 11.73, *p* = .004), and the interaction (*F*_1.3,19.8_ = 7.48, *p* = .008; *df*s are Greenhouse-Geisser adjusted when appropriate). For the category effect, although color and number accuracy did not significantly differ (M = .81 and .86, SD = .05 and .11, respectively; *t*_15_ = 1.86, *p* = .082), they were each higher than ring accuracy (M = .60, SD = .09; respectively, *t*_15_ = 8.55 and 9.13, *p*s < .001). Additionally, whereas the first test probe was associated with higher overall accuracy than the second, the interaction reflected that this order difference was driven by the ring condition (*t*_15_ = 3.67, *p* = .002; colors: *t*_15_ = .34, *p* = .74; numbers: *t*_15_ = .75, *p* = .47).

Response times (RTs) to the recognition probes were faster for correct than incorrect judgments (*t*_15_ = 4.77, *p* < .001), as displayed in Figure 1c. For the remaining RT analyses, we focused on correct judgments as there weren’t enough incorrect judgments in each cell (category × cue/test order) to form reliable means for each subject. The two-way ANOVA revealed only a significant main effect of category (*F*_1.4,20.6_ = 14.52, *p* < .001; see Figure 1d). The pattern of performance was analogous to that found for accuracy, with RTs for colors and numbers not differing significantly (*t*_15_ = 1.70, *p* = .11) but each being faster than responses in the ring condition (respectively, *t*_15_ = 4.14 and 3.60, *p* < .001 and *p* = .0026).

### fMRI results

The fMRI data were analyzed using an MVPA approach (see Haynes & Rees, 2006; Norman et al., 2006; Haxby et al., 2014). Three-way classifications (colors vs. numbers vs. rings) were conducted on an individual-subject basis with logistic regression models of whole-brain patterns (along with feature selection and L2-regularization; see *Whole-brain classification*). To account for the standard delay and expansion of the hemodynamic response, the results reported throughout the main text focus on classifiers trained and tested on patterns from the “peak” period corresponding to 6, 8, and 10 s after the stimulus onset of interest. The corresponding measures for these three time points were averaged and then subjected to statistical testing to mitigate the problem of multiple comparisons associated with testing individual time points. The full time courses and corresponding results of additional comparisons are summarized in each figure and also detailed in Table S1 to provide converging evidence.

Category-related patterns in the FoA

Patterns of brain activity evident during the first presentation array likely provide the “purest” representations in the FoA, given that only one stimulus category is relevant at this point in the trial. To ensure that activity corresponding to the three categories could be effectively distinguished, a classifier was first cross-validated on data associated with this initial array. Figure 2a displays classifier accuracy (M = .47, SD = .12) averaged over all stimulus categories and collapsed across the peak timepoints, which was significantly greater than chance (1/3, given the three categories; *t*_15_ = 4.60, *p* < .001). (As classifier accuracy is expected to exceed the chance level, statistical tests involving this measure are one-tailed unless otherwise noted. Additionally, although directionality becomes more complicated with the switch to measuring classifier evidence, as described below, we continue this approach of using hypothesis-driven one-tailed tests unless noted.) The time course of accuracy for this period, as shown in Figure 2b, additionally showed the expected shape, with four time points (4, 6, 8, and 10 s) in the vicinity of the hemodynamic response peak significantly exceeding the chance level.

**Figure 2.**
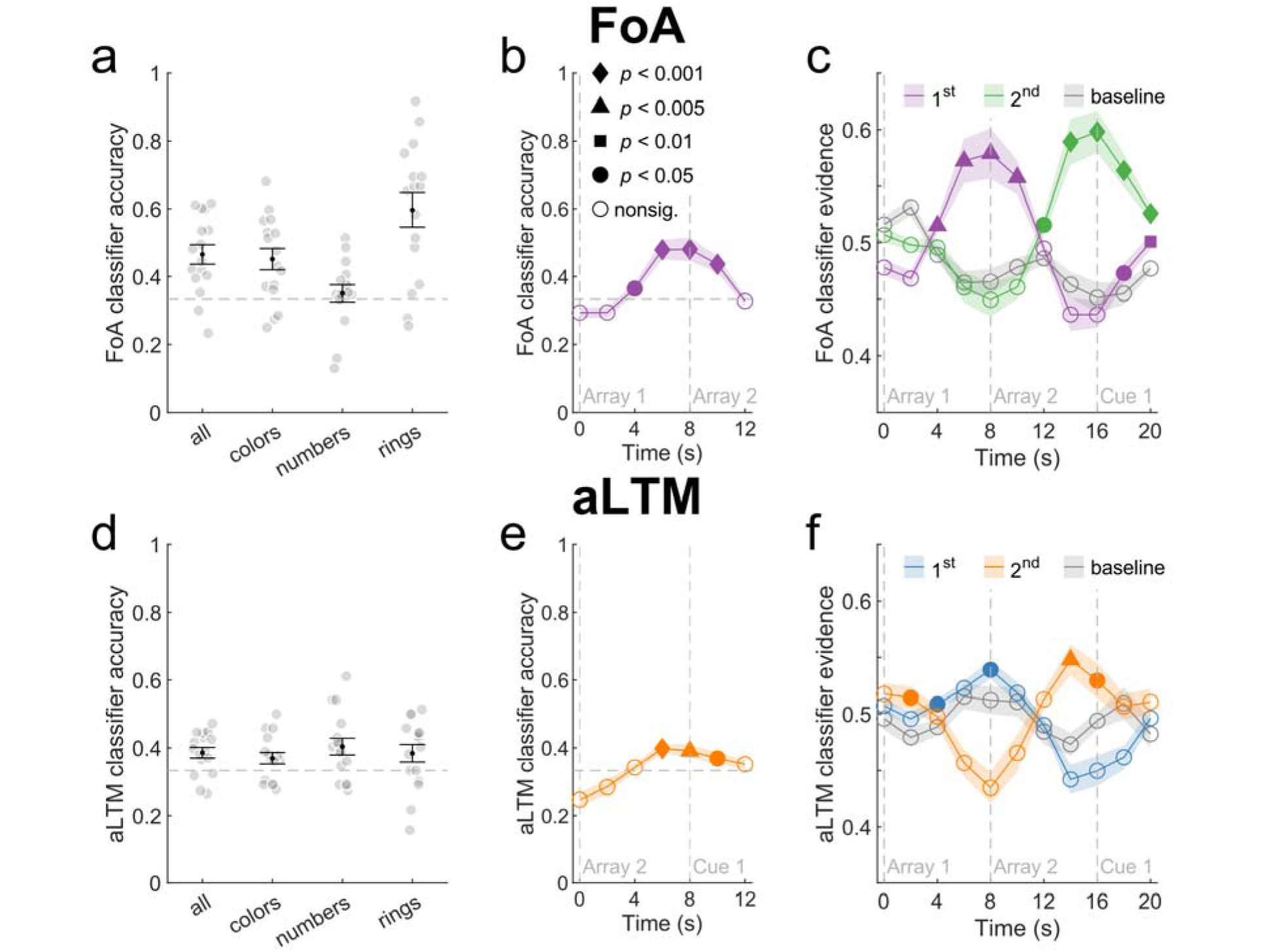
Results of the FoA and aLTM classifiers following the first and second presentation arrays. **(a)** FoA classifier accuracy averaged over the standard peak of the hemodynamic response (6, 8, and 10 s after stimulus onset) following the first presentation array. The results are collapsed over as well as separated according to the three stimulus categories. Black dots are centered on the group means, error bars reflect ±SEM, and transparent dots correspond to individual subject means. The dashed line denotes chance accuracy. **(b)** Time course of FoA classifier accuracy, collapsed over stimulus category (see Figure S3 for separated results). Symbols correspond to the group mean, and the shaded area is equivalent to ±SEM. Dashed lines denote chance accuracy and the onsets of corresponding trial events (Array 1, Array 2). The filled symbols (see legend; also applies to panels c, e, and f) indicate arbitrarily-defined significance thresholds based on one-tailed t-tests vs. chance, respectively corresponding to t-values of 1.75, 2.60, 2.95, and 3.73 (Cohen’s d: .45, .67, .76, and .96). Table S1 of the Supplemental Material provides the exact resulting values at each time point. See the main text for the collapsed results for the peak of the hemodynamic response. **(c)** FoA classifier evidence (as opposed to accuracy) following the presentation arrays. Purple corresponds to the category presented first, green corresponds to the category presented second, and gray corresponds to the category not presented on the trial (baseline). Filled symbols represent significance based on one-tailed t-tests vs. the baseline. **(d)** aLTM classifier accuracy averaged over the standard peak of the hemodynamic response (6, 8, and 10 s after stimulus onset) following the second presentation array. The results are collapsed over as well as separated according to the three stimulus categories. See panel a caption for additional details. **(e)** Time course of aLTM classifier accuracy, collapsed over stimulus category (also see Figure S3). Dashed lines denote chance accuracy and the onsets of trial events (Array 2, Cue 1). **(f)** aLTM classifier evidence following the presentation arrays. Blue corresponds to the category presented first, orange corresponds to the category presented second, and gray corresponds to the baseline category. Filled symbols represent significance based on one-tailed t-tests vs. the baseline.

Accuracy separated for each category is also displayed in Figure 2a and further revealed that classifier performance was primarily driven by the ring condition and, to a lesser extent, the color condition. Accordingly, classifier accuracy differed by category (*F*_1.7,26_ = 18.52, *p* < .001), such that ring accuracy was higher than that for colors (*t*_15_ = 3.60, *p* = .0026), which was in turn higher than that for numbers (*t*_15_ = 3.04, *p* = .0083). Whereas accuracy exceeded chance for the former two conditions (respectively, *t*_15_ = 5.21 and 3.76, both *p*s < .001), it did not for the number condition (*t*_15_ = .70, *p* = .25). (See Figure S3 for the corresponding time courses for each stimulus category.)

### Category-related patterns in aLTM

In contrast with the FoA activity present during the first stimulus array, activity corresponding to aLTM was expected to occur upon presentation of the second array, when trial demands switched to encoding a different category. Cross-validating a classifier for this purpose thus involved labeling the data from the second array with the category present during the first array (as opposed to the category currently being presented). Figure 2d shows overall classifier accuracy, which had a mean of .39 (SD = .06) and was significantly above chance (*t*_15_ = 3.36, *p* = .002). Similar to the FoA results during the first array, accuracy for the aLTM classifier during the second array also followed the expected hemodynamic time course (Figure 2e), exceeding chance from 6-10 s after array onset. Unlike the FoA classifier, however, aLTM accuracy was more balanced across the three stimulus categories (F < 1; see Figure 2d), with each category giving rise to an accuracy level significantly above chance (colors: *t*_15_ = 2.16, *p* = .02; numbers: *t*_15_ = 2.87, *p* = .006; rings: *t*_15_ = 1.96, *p* = .03). (The corresponding time courses of aLTM classification for each category are provided in Figure S3).

As stated in the Introduction, we hypothesized that brain patterns associated with aLTM would be more difficult to identify than those for the FoA. This difference could be due to multiple factors, with the obvious ones being that aLTM activity came from a time in the trial (the second array) that was separated from actual stimulus presentation (the first array) and that it was assessed when there were multiple categories relevant to the current trial. As predicted, overall accuracy for the aLTM classifier was significantly lower than that for the FoA (*t*_15_ = 3.43, *p* = .002) but again above chance in all cases (p < .05).

### Distinguishing the onsets and time courses of FoA- and aLTM-related activity

Having established that category-related brain patterns in the FoA and aLTM could be accurately identified, our analyses from this point forward switched away from accuracy as a measure of classifier performance and to its raw output (or *evidence*) for each of the three stimulus categories (for similar approaches, see Poppenk & Norman, 2012; Leiker & Johnson, 2014; Lewis-Peacock et al., 2015). The motivation for this switch is based on the evidence measure being able to provide separate and continuous indices for each category. In the context of the current classification approach, evidence can range from 0 to 1, with .5 corresponding to evidence neither for nor against a category. Thus, aside from the period of the first stimulus array, when only one relevant category has been presented for the current trial, this measure allows for testing the presence of multiple types of information held in working memory. This is contrary to one of the limitations associated with a binary measure like accuracy which, by nature, requires each time point to be assessed with respect to a category that is deemed “correct”.

Figure 2c displays the evidence measures for the FoA classifier across the two successive stimulus arrays. As shown, the evidence for each presented category peaked in turn, while that for the baseline category not presented on the current trial was relatively stable around .5. Comparing the results for the first-presented category to the baseline (and collapsing over the three stimulus categories to provide more stable measures) revealed a significant enhancement in evidence at multiple time points as well as averaged over the peak period (*t*_15_ = 3.78, *p* < .001). Following FoA evidence through to the second array further revealed the expected switch, with the measure for the second presented category exceeding the baseline (*t*_15_ = 4.73, *p* < .001), but that for the previously presented category no longer exceeding it (*t*_15_ = 1.03, *p* = .84). Directly contrasting the magnitudes of FoA-related evidence across the two arrays, relative to their respective baseline values, revealed that the measure for the second array was significantly higher than that for the first (*t*_15_ = 2.22, *p* = .042, two-tailed).

In contrast to the foregoing classification, Figure 2f displays classifier evidence as trained on the category in aLTM. As with the findings of above-chance accuracy, peak evidence for the category in aLTM exceeded the baseline measure during the second presentation array (*t*_15_ = 2.22, *p* = .021). Additionally, the magnitude of aLTM evidence (subtracting the evidence for the baseline condition) during the second array was smaller than that for the FoA when measured from either the first or second array (*t*_15_ = 2.43 and 3.30, *p* = .014 and .002, respectively). Finally, it is noteworthy that aLTM evidence for a given category dips below baseline while that category is in the FoA. This was evident for both the first (*t*_15_ = 3.23, *p* = .006, two-tailed) and second arrays (*t*_15_ = 2.79, *p* = .014, two-tailed), suggesting that the FoA and aLTM classifiers may be sensitive to distinct patterns of brain activity.

### Retrospective cueing reactivates category-related neural activity for the FoA

A key motivation of the current study was to investigate how retrospective cues elicit reactivation of category-specific neural patterns. As described earlier, the independence of whether each of the two cues targeted the first or second arrays led to two types of trials: repeat and switch (also see Lewis-Peacock et al., 2012, 2015; Rose et al., 2016; Christophel et al., 2018). Figure 3a displays the time courses of FoA activity for repeat trials, where it was apparent that activity for the cued category exceeded the baseline in two separate periods, with the effects for the second compared to the first cueing instance being both delayed (12 vs. 8 s after cue onset) and shorter-lived (two vs. four time points). Additionally, significant differences in peak evidence were present for the first cue (*t*_15_ = 2.47, *p* = .013) but not the second (*t*_15_ = .627, p = .270). For switch trials, the respective cues for the different categories also gave rise to increases in FoA activity, as shown in Figure 3b. Testing the evidence measures during the peak period revealed results similar to those for repeat trials, in the form of reliable differences from baseline only for the first cue (*t*_15_ = 3.19, *p* = .0031; for the second cue, *t*_15_ = 1.55, *p* = .071). As was the case with the repeat trials, though, testing individual time points did reveal significant effects for both cues, albeit again delayed and less sustained for the second.

**Figure 3.**
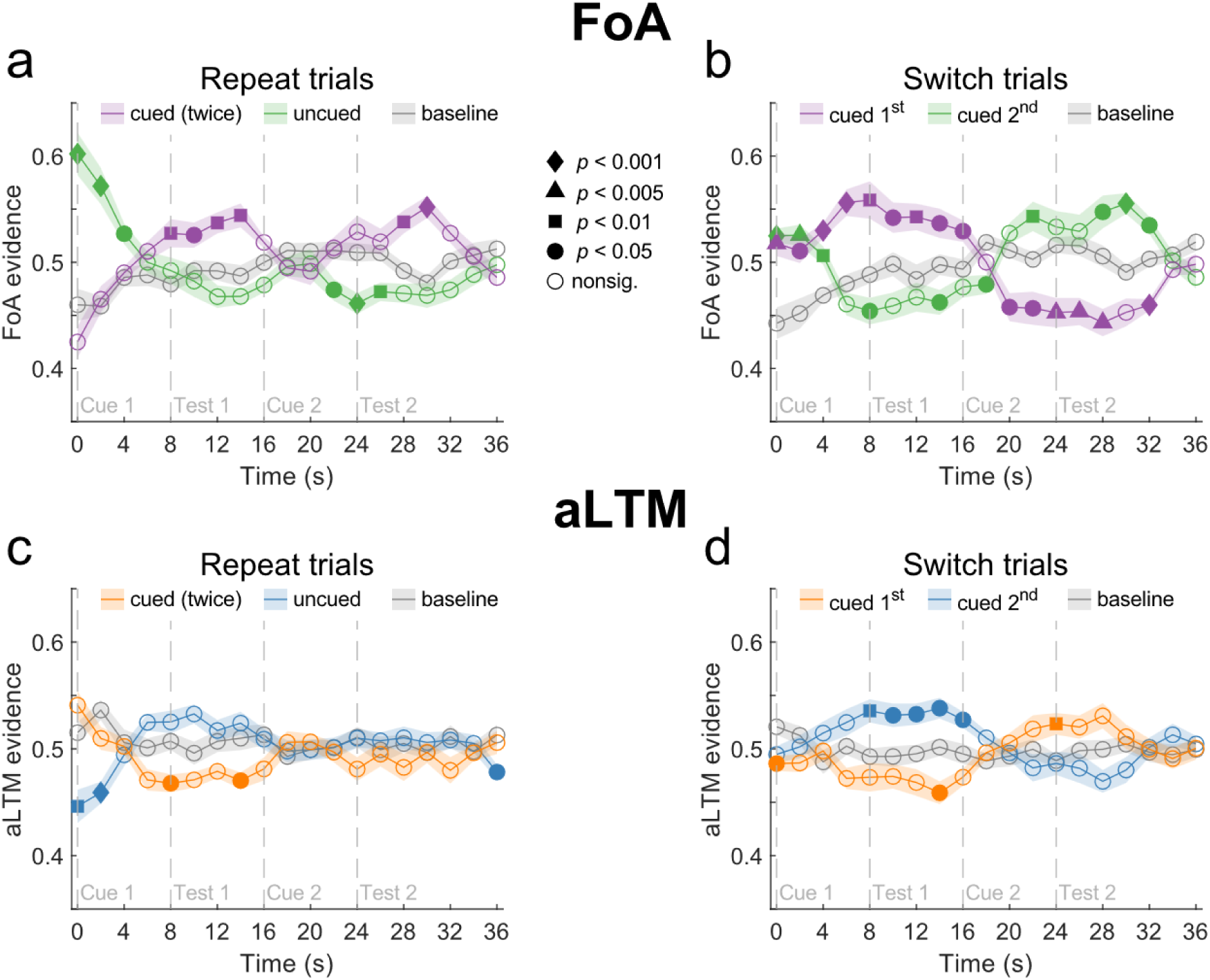
Results of the FoA and aLTM classifiers during the cue and test periods for all repeat and switch trials. **(a)** For repeat trials, time courses of FoA classifier evidence for the twice-cued category (purple), the uncued category (green), and the baseline (gray). Symbols correspond to the group mean, and the shaded area is equivalent to ±SEM. Filled symbols above the baseline indicate different levels of significance based on one-tailed (hypothesis-based) t-tests vs. the baseline, and filled symbols below the baseline indicate significance based on two-tailed (non-hypothesized) tests vs. baseline (also applies to panel b). Dashed lines denote onsets of corresponding trial events. The legend of symbols for significance levels (respective t-values: 1.75, 2.60, 2.95, and 3.73; Cohen’s d: .45, .67, .76, and .96; see Table S1 for exact statistics) applies to all subsequent panels. See the main text for the collapsed results for the peak of the hemodynamic response. **(b)** For switch trials, time courses of FoA classifier evidence (purple, cued first; green, cued second; gray, baseline). **(c)** For repeat trials, time courses of aLTM classifier evidence (orange, cued twice; blue, uncued; gray, baseline). Filled symbols indicate different levels of significance based on two-tailed (non-hypothesized) t-tests vs. the baseline (also applies to panel d). **(d)** For switch trials, time courses of aLTM classifier evidence (orange, cued first; blue, cued second; gray, baseline).

One aspect of cue-elicited reactivation we sought to further address was what happens when a category leaves the FoA but is still potentially relevant to that trial. Our hypotheses about such activity for the uncued category differed for the two cueing instances, and we examine them here in turn. For the first instance, the uncued category was still relevant since it was unknown whether it would be cued later. Yet, it should be temporarily removed from the FoA (Lewis-Peacock et al., 2012). Testing for differences between the uncued and baseline conditions during our standard peak analysis window for this first cueing instance revealed a significant below-baseline (i.e., negative) effect for switch trials (*t*_15_ = 2.77, *p* = .014, two-tailed) but no such effect for repeat trials (*t*_15_ = .29, *p* = .77, two-tailed). When testing individual timepoints, three significant negative effects were evident for switch trials (Figure 3b), but none were evident for repeat trials (Figure 3a; also see Table S1). Additionally, there was increased activity for the uncued condition relative to baseline early on after the first cue; although these results were unexpected, we interpret them as potentially reflecting residual activity from the second stimulus presentation array.

For the second cue, our hypothesis about FoA activity for the uncued category is based on the fact that category-related information is no longer needed for the remainder of the trial. Thus, stronger below-baseline effects could signify the removal of irrelevant information from working memory. Indeed, such effects were evident during the peak period for both repeat and switch trials (respectively, *t*_15_ = 4.58 and 4.11, *p*s < .001, two-tailed), with multiple individual time points also exhibiting significance for each trial type (see Figures 3a and 3b).

### Cueing elicits opposing FoA and aLTM effects for the same category

We next sought to dissociate the activity patterns associated with the FoA vs. aLTM, dependent on which stimulus category is currently being cued. Figures 3c and 3d provide the analogous time courses of aLTM activity for repeat and switch trials. As shown and expected on the basis of the findings during the array periods, aLTM activity appeared weaker in magnitude than that for the FoA. More importantly, though, the pattern of aLTM activity for the cued and uncued categories took a form that was opposite of the FoA. In particular, for both repeat and switch trials, aLTM activity for the category cued first was significantly below the baseline level at multiple time points (see Table S1 for details), with these effects for the peak period reaching significance only on repeat trials (*t*_15_ = 2.63, *p* = .019, two-tailed; for switch trials, *t*_15_ = 1.69, *p* = .11). By contrast, aLTM activity for the uncued category during the first cue-and-test period was at least numerically greater than the baseline, with multiple time points (8 to 16 s after cue onset) and peak evidence being significant for switch trials (*t*_15_ = 2.74, *p* = .015, two-tailed; for repeat trials, *t*_15_ = 2.12, *p* = .051). Given these somewhat inconsistent results for aLTM, and the fact that the below-baseline effects are novel, we further test the dissociation of FoA and aLTM activity using a more optimal set of trials described below.

The optimal data needed to test this dissociation follows the first cue to attend and is from only those trials in which the first-cued category was presented in the first array, since there should be both FoA and aLTM representations of that category that can be compared (whereas the category presented in the second array, when cued first, might only exist in the FoA form). Figures 4a and 4b display the time courses of FoA and aLTM activity during this period, along with their corresponding baseline conditions. FoA activity was significantly greater than baseline (*t*_15_ = 3.49, *p* = .0016), but the opposite effect was observed for the aLTM representation of the same cued category (*t*_15_ = 3.07, *p* =.0039), with both of the effects sustained over a long time period (also see Table S1).

**Figure 4.**
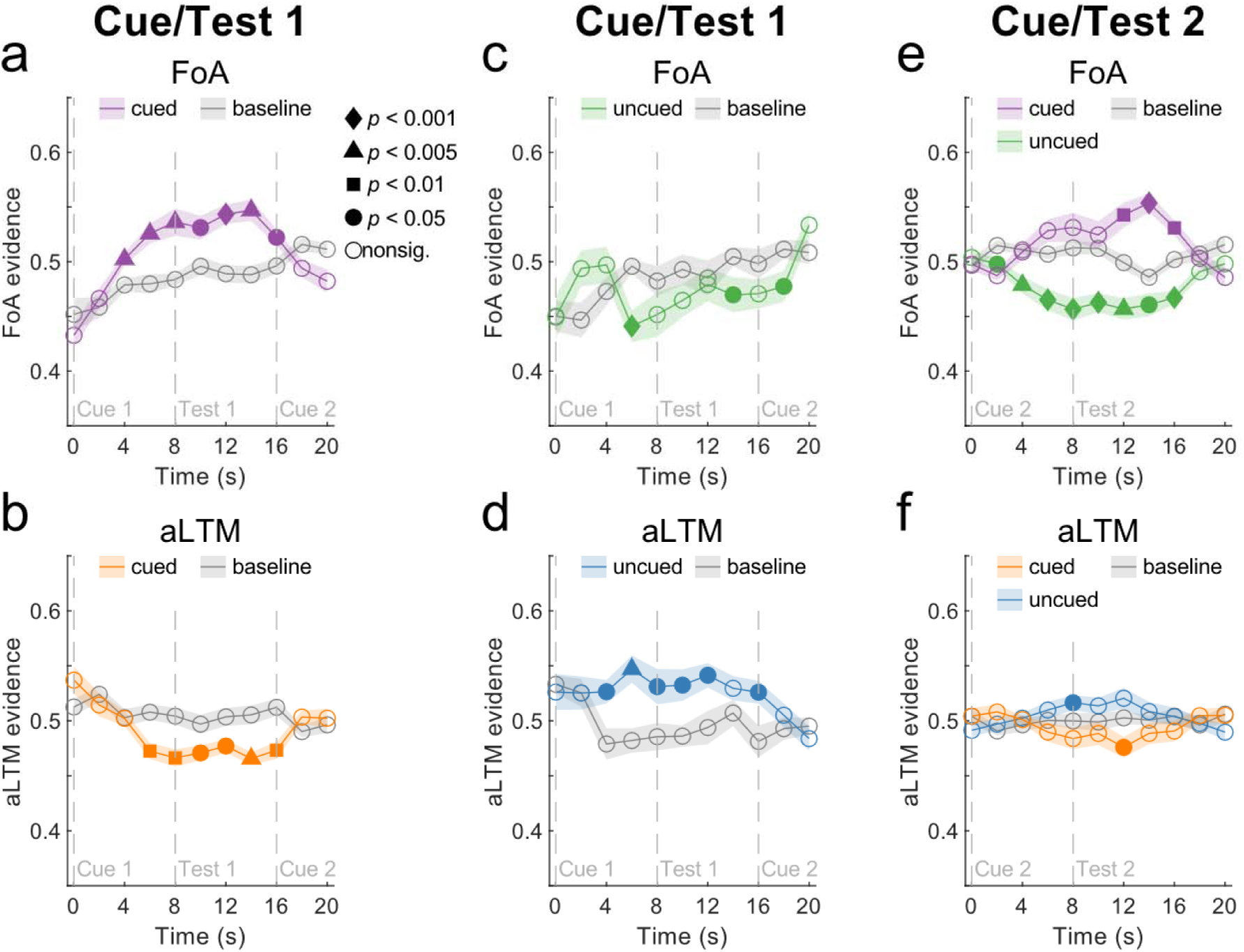
Results of the FoA and aLTM classifiers during the cue and test periods, restricted to “optimal” trials where both states exist for the same stimulus category. **(a, b)** Time courses of FoA (panel a) and aLTM (b) classifier evidence for the cued category during the first cue/test period, along with the respective time courses for the baseline category. Note that these data correspond only to “optimal” trials, which, as discussed, are trials in which the first-cued category was from the first array, producing both FoA and aLTM measures for the same category. Symbols correspond to the group mean, and the shaded area is equivalent to ±SEM. Filled symbols indicate significance levels based on one-tailed (hypothesis-based) t-tests above (for FoA) and below (for aLTM) the baseline. The legend of symbols for significance levels (respective t-values: 1.75, 2.60, 2.95, and 3.73; Cohen’s d: .45, .67, .76, and .96; see Table S1 for exact statistics) applies to all subsequent panels. (See Figure S4 for corresponding results separated by stimulus category.) **(c, d)** Time courses of FoA (c) and aLTM (d) classifier evidence for the uncued category during the first cue/test period, along with the respective baseline. As in panels a and b, these data correspond only to trials in which both FoA and aLTM representations are present for a category on the same trial. **(e, f)** Time courses of FoA (e) and aLTM (f) classifier evidence during the second cue/test period.

As was demonstrated for the presentation array period, the FoA classifier was primarily driven by the ring category, with the number category failing to give rise to above-chance accuracy (see Figure 2a and Figure S3). One possible explanation for this difference is that the visually-presented numbers can be readily translated into their verbal labels, whereas exemplars within the color and ring orientation categories might be more easily confused. To further test whether numbers are possibly held in WM in a manner distinct from that for the other categories, whereby aLTM is relied upon more and involvement of the FoA is minimized, we additionally examined cue-related evidence for the two classifiers separately for each stimulus category. Figure S4 shows that, whereas FoA evidence was again strongest for the ring category, there was no indication of enhanced aLTM activity for numbers (i.e., all three categories instead showed the below-baseline effect described above).

The complementary effect for the cued category described above is what is not attended to during the first cue period. For the uncued category, the prediction is that aLTM activity should increase, given the possibility that this category will be relevant during the second period. Further, given the opposing effects shown above, FoA activity for the uncued category should as a result be diminished. Figures 4c and 4d display these time courses. Whereby aLTM activity for the uncued category was significantly greater than the corresponding baseline (*t*_15_ = 3.34, *p* = .0022), FoA activity for the uncued category was below baseline (*t*_15_ = 2.58, *p* = .0104). This pattern is consistent with aLTM keeping the uncued category available for later use during the trial, but at the expense of its FoA representation.

### Residual evidence of task-irrelevant representations

As noted above, category-related activity in the FoA was not as prominent upon presentation of the second cue as it was following the first. Figure 4e shows FoA activity elicited by the second cue but now collapsed over repeat and switch trials. Significance for the cued category was met for three individual time points (12-16 s) but it did not reliably differ from baseline in the earlier, peak interval (*t*_15_ = 1.20, *p* = .124). Additionally, FoA activity for the uncued category was below the baseline level for both the peak period (*t*_15_ = 6.49, *p* < .001) and at several individual time points (from 2 to 16 s).

One final aspect about the second cue period that is unknown concerns what happens to aLTM representations once they are no longer needed for the trial. As was demonstrated for the first cue, aLTM activity for the cued category was below that for the baseline category, consistent with making way for accessing the stronger FoA representation. Likewise, aLTM activity for the uncued category during that first interval exhibited enhancement, presumably allowing category information to be maintained for later use in the trial. A straightforward prediction based on the latter result is that aLTM activity for the uncued category during the second cue period should be indistinguishable from the baseline, given that it is no longer needed. Alternatively, it is possible that residual aLTM activity is detectable before it fully decays. As shown in Figure 4f, aLTM activity did differ from the baseline condition. Analysis of the peak period revealed a significant enhancement for the uncued category relative to baseline (*t*_15_ = 2.09, *p* = .027), but the effect for the cued category was not reliable (*t*_15_ = .92, *p* = .187). When examining individual time points, there was further evidence that the uncued aLTM time course was above baseline (at 8 s) and some evidence of a cued effect below baseline (at 12 s).

### Topographical similarities and differences in FoA- and aLTM-related activity

Given the opposing time courses of FoA and aLTM activity, the final aspect of these two states we sought to address was the nature of any differences with respect to their brain topographies. There are multiple possibilities for such differences, including representation by disparate brain regions, distinct representations in a common set of regions, or a combination of both.

To understand the activity patterns, we first examined the voxel-wise *importance* values resulting from training the FoA and aLTM classifiers. Importance corresponds to the product of a voxel’s mean activity during the presentation of each stimulus category array (i.e., the training period, which is the first array for FoA and the second array for aLTM) and its trained classifier weight for each category. The resulting importance maps for the two classifiers are respectively displayed in Figure 5. As described in *Searchlight classification*, these maps are purely for descriptive (i.e., not statistical) purposes, as they are averaged across subjects and arbitrarily thresholded (≥|.0001|). Most of the resulting voxels were in lateral and medial parietal and lateral prefrontal cortices. As is apparent from the figure, the FoA maps (Figures 5a and 5b) were more extensive than the aLTM maps (Figures 5c and 5d), with 2,178 vs. 509 voxels respectively passing the pre-determined threshold. Additionally, and again on a purely descriptive basis, overlapping voxels across multiple stimulus categories were largely restricted to the FoA classifier and were evident most prominently in the vicinity of bilateral intraparietal sulcus (IPS; see Figure 5b). Finally, given that the FoA classifier was largely driven by the ring category (see Figures S3 and S4), it is also not surprising that most of the suprathreshold voxels are from that condition (83% vs. 10% and 7% for the color and number conditions, respectively). This imbalance was less pronounced but still present for the aLTM classifier (47, 28, and 26% for the ring, color, and number conditions).

**Figure 5.**
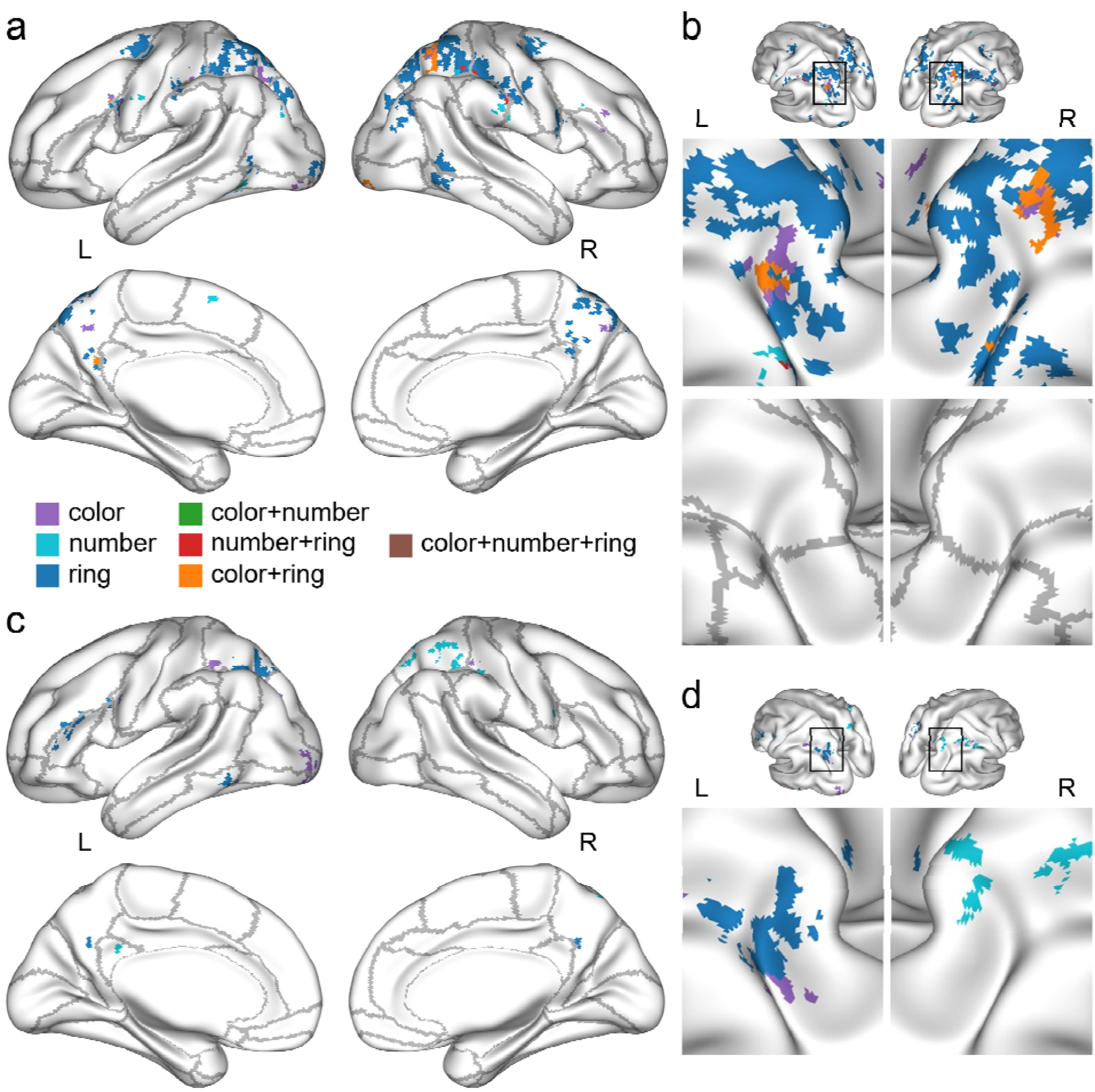
Descriptive brain maps of “important” voxels from the FoA and aLTM classifiers, for each stimulus category and their overlap. **(a)** Voxel-wise importance values (activity×weight) from the trained FoA classifiers, after averaging across subjects and extent-thresholding (*k* ≥ 10). Voxels are color-coded according to being unique to each stimulus category as well as overlapping among combinations of categories (see legend). **(b)** Voxel-wise importance values for the FoA classifiers, as shown in panel a, highlighting the overlap among stimulus categories in the vicinity of intraparietal sulcus (IPS). Outlined boxes at the top indicate the zoomed-in portion (middle). Anatomical (AAL) boundaries are shown in gray separately at the bottom so as not to occlude the results. All maps are rotated at 45° elevation and ±45° azimuth. (See panel a for color legend.) **(c)** Voxel-wise importance values (activity×weight) from the trained aLTM classifiers, after averaging across subjects and extent-thresholding (*k* ≥ 10). (See panel a for color legend.) **(d)** Voxel-wise importance values for the aLTM classifiers, as shown in panel c, highlighting the IPS. Note that there was no overlap among stimulus categories, in contrast to the FoA overlap shown in panel b. Outlined boxes at the top indicate the zoomed-in portion (bottom). All maps are rotated at 45° elevation and ±45° azimuth. (See panel a for color legend and panel b for AAL boundaries.)

To further address any differences between the FoA and aLTM patterns in a statistical manner, we conducted searchlight analyses that involved passing a small sphere (of 33 voxels) throughout the brain to classify category-related activity. The resulting maps of searchlights exhibiting significant above-chance accuracy, as displayed in Figure 6, revealed many of the same regions as the importance maps did (see Figure 5), mostly including bilateral and medial parietal cortex, for both the FoA and aLTM classifications (see Table S2 for the full list of significant clusters). In contrast to the limitations of assessing overlap with the importance maps, though, the searchlight analyses allowed for statistically testing for overlap (based on an inclusive-masking procedure using a conjoint threshold of *p* < .001 and *k* ≥ 10 voxels; also see Table S3). As shown in Figure 6c, the results for the two classifications exhibited significant overlap along the IPS. Additionally, statistically contrasting the searchlight maps for the FoA and aLTM classifications revealed greater activity for the FoA in multiple regions, again mostly in the vicinity of bilateral superior occipital and parietal cortices, as displayed in Figure 6d (see Table S3). There were no significant differences, however, in the opposite direction.

**Figure 6.**
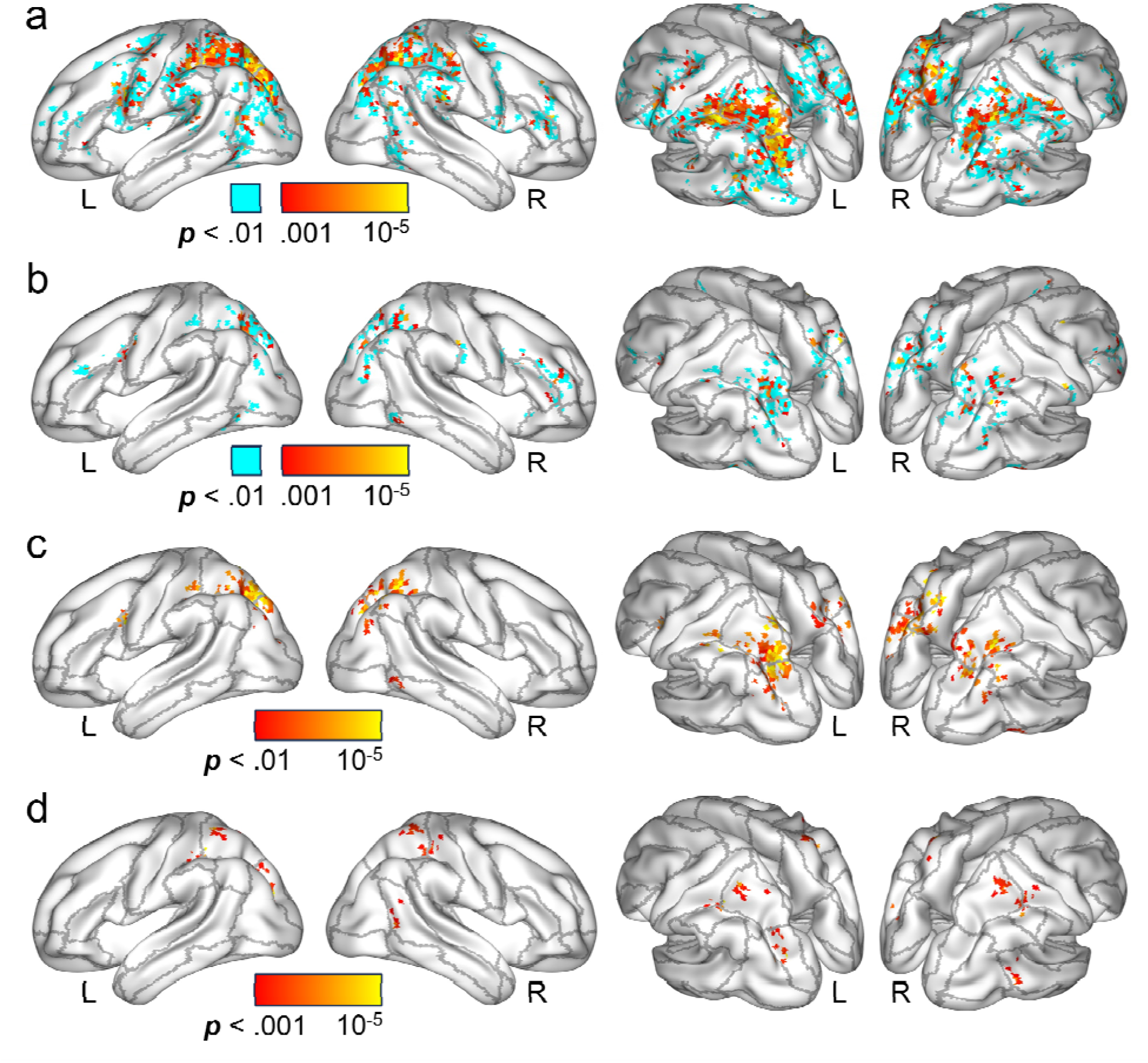
Statistical brain maps of searchlight classification results for FoA and aLTM, along with their overlap and differences. **(a)** Significant results (in hot colors; *t*_15_ > 3.73, *p* < .001, *k* ≥ 10 voxels; *p* < .05, FWE-corrected) from the searchlight analyses of FoA-related accuracy. Voxels not significant by our *p* < .001 criterion are also highlighted (in cyan; *t*_15_ > 2.60, *p* < .01, *k* ≥ 10 voxels), as they are potential results for the inclusive-masking procedure (see panel c). The maps on the right are rotated at 45° elevation and ±45° azimuth to highlight the results in the vicinity of intraparietal sulcus (IPS; this also applies to the subsequent panels). Anatomical (AAL) boundaries are shown in gray. The full medial views are not displayed here, as significant medial voxels were mostly posterior and visible on the rotated surfaces. Also see Table S2 for details of significant clusters. **(b)** Results from the searchlight analyses of aLTM-related accuracy. (See panel a caption and Table S2 for further details.) **(c)** Significant overlapping results from the FoA and aLTM searchlight analyses. Each of the two contrasts was thresholded at *p* < .01 and then submitted to an inclusive-masking procedure, resulting in a final, conjoint threshold of *p* < .001 (*k* ≥ 10 voxels). (See panel a caption and Table S3 for further details.) **(d)** Results of greater searchlight-based accuracy for FoA than aLTM (*p* < .001, *k* ≥ 10). No significant clusters were evident for the reverse contrast (aLTM > FoA). Also see Table S3.

To facilitate comparison between the current results and those of Christophel et al. (2018), which served as motivation for multiple aspects of our design and analysis approach, we collapsed the aforementioned searchlight outcomes into the regions of interest (ROIs) they employed (based on Wang et al., 2015). Consistent with their ROI results and our higher-resolution searchlight results, intraparietal sulcus (IPS; IPS0-IPS5 of Wang et al.) exhibited above-chance accuracy for both the FoA and aLTM classifications (respectively, *t*_15_ = 5.75 and 5.43, *p*s < .001). Moreover, we again observed significantly greater activity for the FoA in this region (*t*_15_ = 4.70, *p* < .001; also see Figure 6c), whereas Christophel et al. did not. For the FEF, a similar pattern emerged, with the FoA and aLTM classifications both producing significant above-chance accuracy (respectively, *t*_15_ = 3.89 and 4.46, *p*s < .001) and the former exceeding the latter (*t*_15_ = 2.53, *p* = .023). Note that this finding of FoA>aLTM activity in frontal cortex was not evident from the searchlight analyses alone, suggesting that averaging the results into ROIs could have contributed to increased statistical power. Finally, Christophel et al. reported significant FoA activity for occipital cortex (OC; V1-V4 of Wang et al.), but we found that the results collapsed into this ROI failed to reach significance for both the FoA and aLTM classifications (respectively, *t*_15_ = 1.05 and .28, *p*s = .156 and .391). Although these discrepant findings for OC might indicate something theoretically important that differs between the two studies, it is also possible that their classification of orientation stimuli was carried out in lower visual areas than our classification of multiple stimulus categories.

## DISCUSSION

Despite consistent findings of neural activity associated with information held in an attended state (focus of attention, FoA) of working memory (Lewis-Peacock et al., 2012; Rose et al., 2016), evidence for a neural correlate of information outside the FoA, and instead in a state of activated long-term memory (aLTM), is inconclusive. Here, we used a retrospective cueing task in combination with MVPA of fMRI to train a classifier on information in the FoA, corresponding to stimuli just presented, and also to train a separate classifier on information in aLTM, based on stimuli that had been presented earlier in a trial but had left the FoA. Our primary findings indicated active neural representations of categorical stimuli that were not relevant to the task goals of the present moment but could potentially be relevant later in the trial. These non-silent neural signatures contrast with the activity-silent form that has been proposed to be used in prior studies (for reviews, see Stokes, 2015; Barbosa et al., 2020) and extend recent findings that were restricted to single stimulus categories, such as visually-oriented gradients (Christophel et al., 2018) and naturalistic images (Paluch et al., 2025), into the domain of multiple categories of information.

Relative to neural activity corresponding to stimuli in the FoA, aLTM-related activity was notably weaker, particularly related to the levels of classifier accuracy and evidence. This could be due to the simple fact that the classifier for the latter state was trained on categories from the previous array (∼8 s earlier) and was thus temporally removed from presentation of the relevant stimuli. Additionally, given that stimuli from other categories were being presented during this second array, they would have activated distinct patterns related to visual stimulation, thus potentially compromising the classification in terms of differences at the level of representation or anatomy (i.e., the classification is less driven by sensory regions). Consistent with this quantitative difference, we also observed that the extent of brain regions exhibiting aLTM-related activity was smaller than that for the FoA (see Figures 5 and 6). Moreover, considerable portions of the voxels identified with the aLTM classifier were a subset of those identified for the FoA (43 and 59%, respectively, based on whole-brain importance values and searchlight classifications). Despite the differences in strength and extents of these two effects, we emphasize that aLTM activity is unlikely to reflect “residual” FoA activity. Indeed, FoA activity for the first array returns to baseline after it peaks, while aLTM activity for the second array begins below baseline (compare the upper and lower panels of Figure 2); evidence of residual activity would instead take the form of a continuation across these time periods. (To entirely rule out this alternative explanation, future studies could employ longer, and perhaps temporally jittered, inter-array intervals.) Together, these findings suggest that the differences in FoA and aLTM could be driven either by activity in the nearby but non-overlapping regions of bilateral superior parietal cortex that are associated with the FoA or by distinct patterns of representation in the common set of voxels.

A novel aspect of the current findings is related to the relationship between the two states of working memory. As hypothesized, our task allowed evidence of FoA- and aLTM-related neural representations to be tracked over the same time periods, whether they were elicited by the task cues (or were uncued) or existed simultaneously for the same stimulus category. Specifically, whereas cueing a given category gave rise to prominent activation of its FoA representation, the aLTM representation for the uncued category was also activated, consistent with it being offloaded to aLTM for potential access later in the trial. Though our intention was to ensure that each category had both FoA and aLTM representations (the former from when it was presented and the latter from when the presentation array changed to a new category), a noteworthy aspect of our task—that the two presentation arrays for a trial were non-orthogonal (i.e., they always drew from two different categories)—also allows for an alternative interpretation of the results. Namely, assigning activity during the second array to the previously-presented category could lead to training the later classifier on the absence of the first-presented category (or “anti-category”) rather than on an additional memory state. As a result, it would be unsurprising that the second classifier, given that it is based on categorical information that is a mixture of the other two categories being presented, would exhibit weaker activity than one trained on the first array. It does not seem possible, at least to us, to distinguish these two accounts with the current data set. While we interpret the second (aLTM) state as providing later access to information that needs to leave the FoA, the alternative interpretation might reflect an attempt to suppress prior information in order to free up attentional resources for the second array. Such suppression must also be specific enough so as not to include the third (baseline) category that is not present on the current trial. Nevertheless, follow-up studies could begin to address this issue by implementing catch trials where the same category is sometimes presented twice, or by manipulating the similarity of stimulus categories, with the expectation that similar stimuli presented in succession might encourage greater use of the anti-category strategy. Despite the nature of the second categorical representation being unclear, the findings reported here provide novel evidence indicating opposing effects for multiple forms of representation, whereby increased activity for one is evident in the face of significant deactivation for the other.

In addition to the opposing time courses of FoA and aLTM representations, another interesting feature distinguishing the patterns for these two memory states was evident during the second cue-and-test period, when only a single category remained relevant. Consistent with what was demonstrated for the first cue period, the aLTM representation for the cued category during the second period appeared to give way to the corresponding FoA representation for that category. One possible interpretation of this opposition is that the stronger (FoA) representation appears to be targeted for use by the subject at the expense of the weaker (aLTM) one. A further prediction relevant to the second cueing period, though, is that activity related to the uncued category will return to baseline. Although the size of the effect was somewhat small, the current results instead indicated that aLTM-related activity for the uncued category remained significantly above the baseline level. This residual enhancement thus provides novel evidence that aLTM patterns might be subject to gradual decay (see Rose et al., 2016), in keeping with some behavioral results (e.g., Ricker et al., 2020). Moreover, these findings highlight a contrast with the FoA effects for cued and uncued categories at this late stage of the trial which, respectively, still appear to be under the influence of rapid (and flexible) activation and inhibitory control. An important aim for future research should be to further characterize the longer-term temporal dynamics of FoA and aLTM neural representations, particularly as they relate to trace durability and decay.

As described in the Introduction, Christophel et al. (2018) demonstrated activity related to holding visual orientation stimuli in a non-FoA state that allowed stimuli to be returned to after initially serving in an uncued condition. The evidence of aLTM neural patterns provided by the current study is consistent with their findings. Although there are several differences between these two studies, the most notable among them is the use of different exemplars of one visual category versus stimuli from multiple categories. The current findings thus provide an important extension to the previous work. That is, the use of a single category, as was the case with Christophel et al.’s oriented gradients, could involve holding the difference between two stimuli in working memory (Gossaries et al., 2018; Cai et al., 2020), as opposed to offloading uncued stimuli from a primary to secondary state to make way for cued stimuli. With multiple categories (also see Lewis-Peacock et al., 2012; Rose et al., 2016), though, capturing the differences between particular categories (or stimuli) likely becomes much more challenging or impossible. The multi-category approach, therefore, can crucially emphasize the functional role of simultaneously using multiple representational states.

It is also important to point out that different categories might not map onto the FoA and aLTM states in a similar manner. Notably, evidence aligning with this idea was demonstrated in multiple forms here. First, the ring category demonstrated the largest FoA effect while the number category resulted, numerically, in the largest aLTM effect (but no evidence of an FoA effect). Although this discrepancy was not predicted, in hindsight it makes sense given that subjects might need to exert increased effort and spend additional time maintaining the relatively novel stimuli of the ring category. By contrast, subjects presumably have extensive experience maintaining number stimuli, and thus the corresponding brain patterns might be weaker or more easily transferred from the FoA to aLTM. With the categories employed here (or with any distinct categories, for that matter), it is challenging to entirely rule out the idiosyncratic differences between stimuli that could correspond to difficulty/effort, familiarity/fluency, verbalization, or even eye movements. Second, the mapping of important voxels associated with each stimulus category exhibited differences across the FoA and aLTM classifications (e.g., compare the voxels highlighted for the color category in Figure 5b versus 5d). These findings could reflect the fact that, whereas some stimuli (e.g., ring orientations) might be best maintained in a perceptual (stimulus-driven) format across memory states, other stimuli (e.g., numbers) might be recoded from the perceptual (visual) level to either an alternative (e.g., phonological) or deeper conceptual level (also see Rose, Craik, & Buchsbaum, 2015; Yue et al., 2019). Such recoding away from the stimulus-driven format is consistent with the classification results being strongest in higher, parietal regions (Rademaker, Chunharas, & Serences, 2019) as opposed to also including lower, perceptual areas (as in Christophel et al., 2018; also see Iamshchinina et al., 2021).

In summary, the findings reported here, in combination with a handful of other studies utilizing pattern classification of fMRI and within-trial retrospective cuing, provide further neural evidence that working memory includes a state beyond the FoA, consistent with the concept of aLTM (Cowan, 1988, 2019; Cowan et al., 2024). More specifically, the current study characterizes multiple features of aLTM representations that contrast with those related to the FoA, such that they can be identified (trained) separately from background activity, exhibit simultaneous but opposing effects when cued, and seemingly decay at different rates. Together with recent work on visual working memory, there is growing and compelling support that this secondary state of working memory constitutes active neural patterns in addition to the activity-silent representations previously proposed (Kamiński & Rutishauser, 2020; Stokes, 2015).

## ACKNOWLEDGEMENTS

We thank Bret Glass for assistance with MRI acquisition. This research was supported by funding from the Department of Psychological Sciences, University of Missouri.

## CRediT AUTHORSHIP CONTRIBUTION STATEMENT

**J. Johnson:** Conceptualization, Formal analysis, Investigation, Methodology, Visualization, Writing – original draft, Writing – review and editing. **S. Christ:** Conceptualization, Methodology, Visualization, Writing – original draft, Writing – review and editing. **N. Cowan:** Conceptualization, Methodology, Visualization, Writing – original draft, Writing – review and editing.

## Notes

https://osf.io/uanqc

### Competing Interest Statement

The authors have declared no competing interest.

