## Supplemental Material for "Active Neural Representation of Stimulus Categories Outside of the Focus of Attention"

**Figure S1. Accuracy results of additional classifiers following the first (for FoA) and second (for aLTM) presentation arrays**

To test whether our pre-experimental choices of classifier (L2-regularized logistic regression) and classifier parameters (penalty = 100; 10,000 features with largest F-values across conditions) did not unexpectedly lead to overfitting, a series of post-hoc analyses were conducted. Accuracy in each of the plots below is plotted for the FoA (in purple) following presentation of Array 1 and for aLTM (in orange) following presentation of Array 2. Accuracy corresponds to the traditional “peak” of the hemodynamic response, averaged over 6, 8, and 10 secs after array onset. For comparison, accuracy for the FoA and aLTM classifiers reported in the main text was .47 and .39, respectively, and comparable to the results shown here, suggesting that our reported results did not lead to increased overfitting.

(a, b) Accuracy from L1- and L2-regularized logistic regression, based on the top 10,000 voxels (with largest F-values) and according to regularization penalties ranging from 100 to 7,500.

(c) Accuracy from L2-regularized logistic regression, based on a liberal regularization penalty of 75% of voxels (e.g., 75 for 100 voxels, and 7,500 for 10,000 voxels) and according to different numbers of voxels (ranging from 100 to 10,000, with the largest F-values).

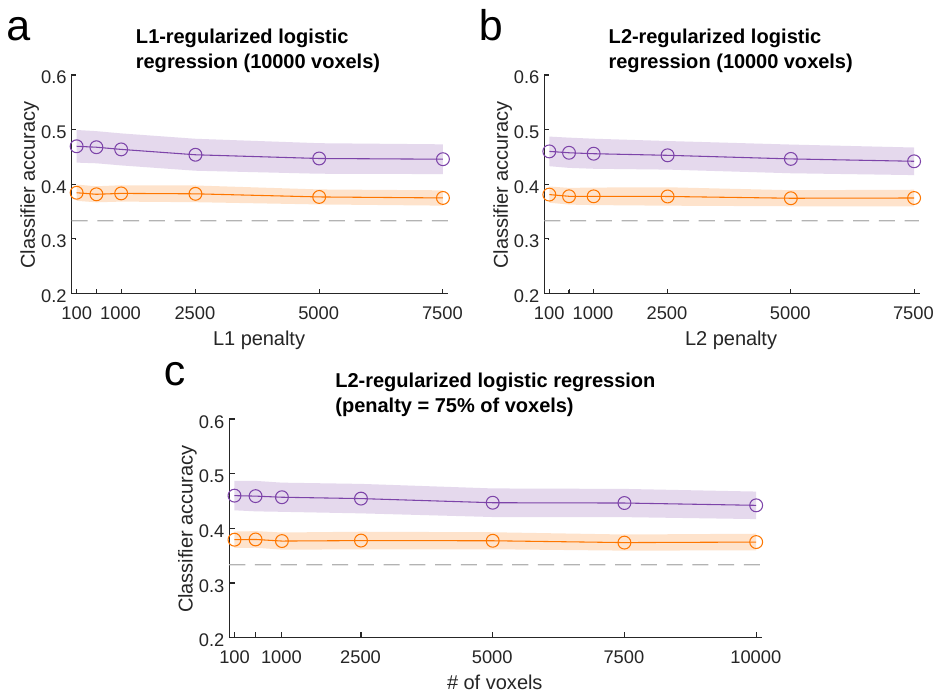

**Figure S2. Behavioral results according to stimulus category and presentation array**

An additional manipulation in the behavioral procedure was whether the cued and tested stimuli were presented earlier in the trial in the first vs. second arrays. Correct response percentages and correct RTs are summarized respectively in panels a and b of the figure below, separated according to stimulus category and whether encoding of that category occurred during the first vs. second array (labeled 1 vs. 2).

Though there were not enough trials to fully explore the three-way interaction (category × array order × cue/test order), we briefly report the results of analyzing the former two manipulations together. Two-way ANOVAs revealed no main effect of array order for correct percentages or associated RTs (respectively, *F*_1,15_ = 3.08 and 1.47, *p* = .1 and .25), as well as no interactions (both *F*s < 1). Consistent with behavioral results reported in the main text (also see Figure 1), the category main effects remained significant for both measures (respectively, *F*_1.8,27.6_ = 50.71, *p* < .001, and *F*_1.3,18.8_ = 9.53, *p* = .004). Black dots are centered on the group means, error bars reflect ±SEM, and transparent dots correspond to individual subject means.

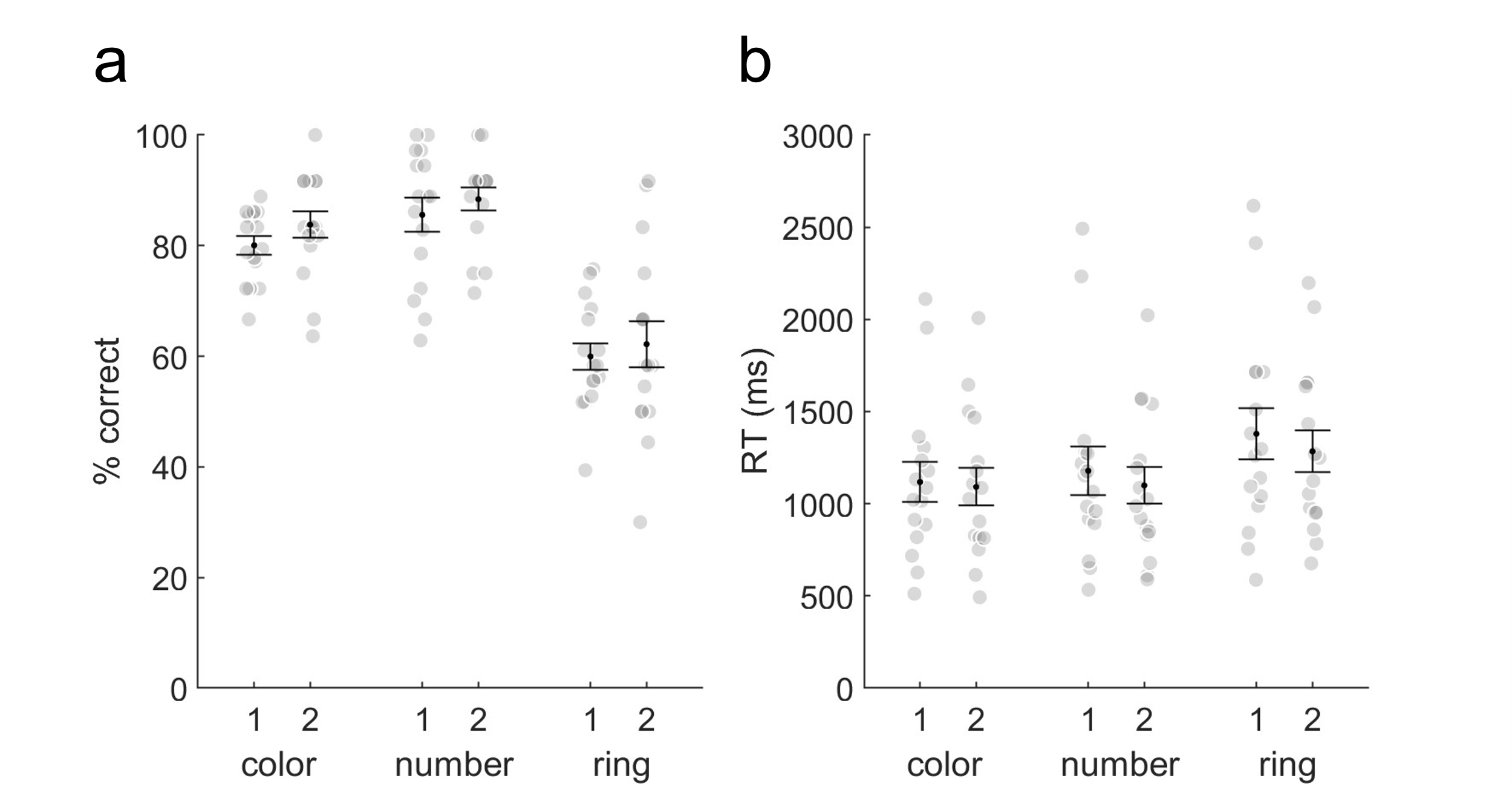

**Figure S3. Classifier accuracy according to each stimulus category**

Proportions of each category selected by the classifier according to the actual condition, for FoA (panel a) and aLTM (panel b). (Chance performance would correspond to .33 in each cell of a given column.) Time courses of classifier accuracy are also provided for FoA (panels c‒e) and aLTM (panels f‒h), separated by stimulus category. Symbols correspond to the group mean, shaded area is equivalent to ±SEM, and filled symbols indicate different levels of significance based on a one-tailed t-tests vs. chance (see legend in panel a). The dashed lines denote chance accuracy and trial events (A1/A2, first/second presentation array; C1, first cue to attend).

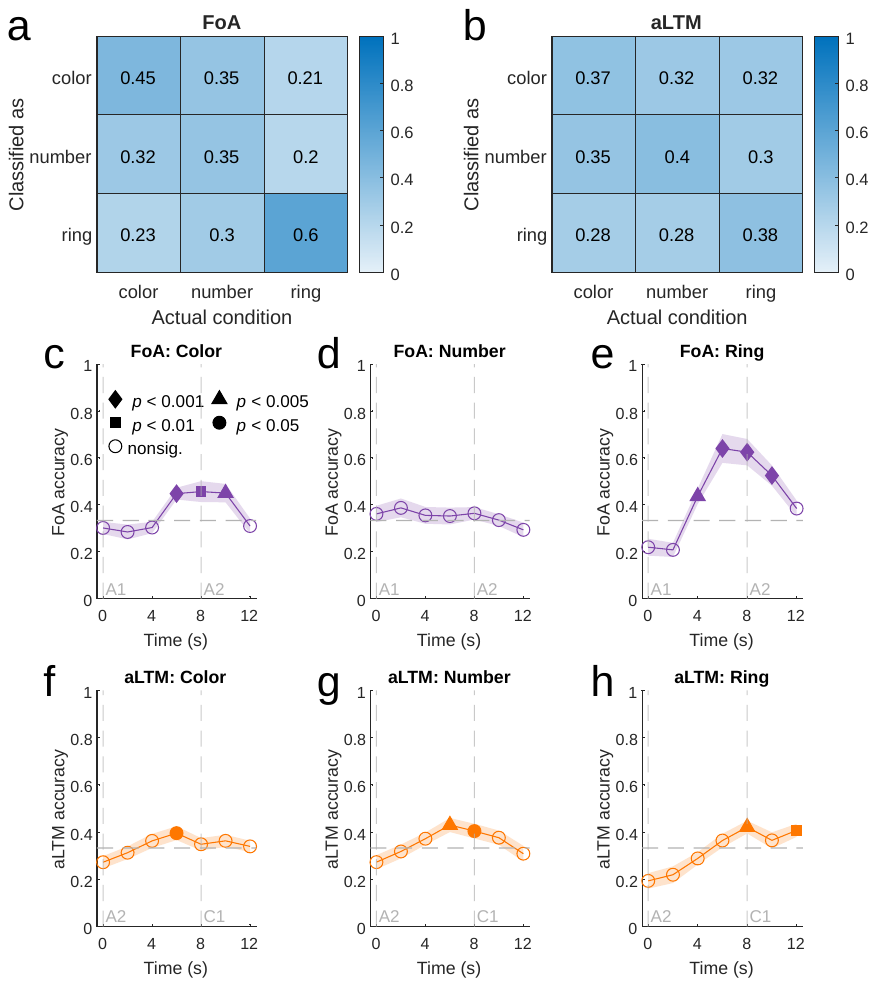

**Figure S4. Classifier evidence during the first cue and test period according to each stimulus category**

Time courses of classifier evidence are provided for FoA (panels a‒c) and aLTM (panels d‒f), separated by stimulus category. Symbols correspond to the group mean and the shaded area is equivalent to ±SEM. Filled symbols (see legend in Figure S3c) indicate different levels of significance based on one-tailed (hypothesis-based) t-tests above (for FoA) and below (for aLTM) the baseline. Dashed lines denote onsets of corresponding trial events.

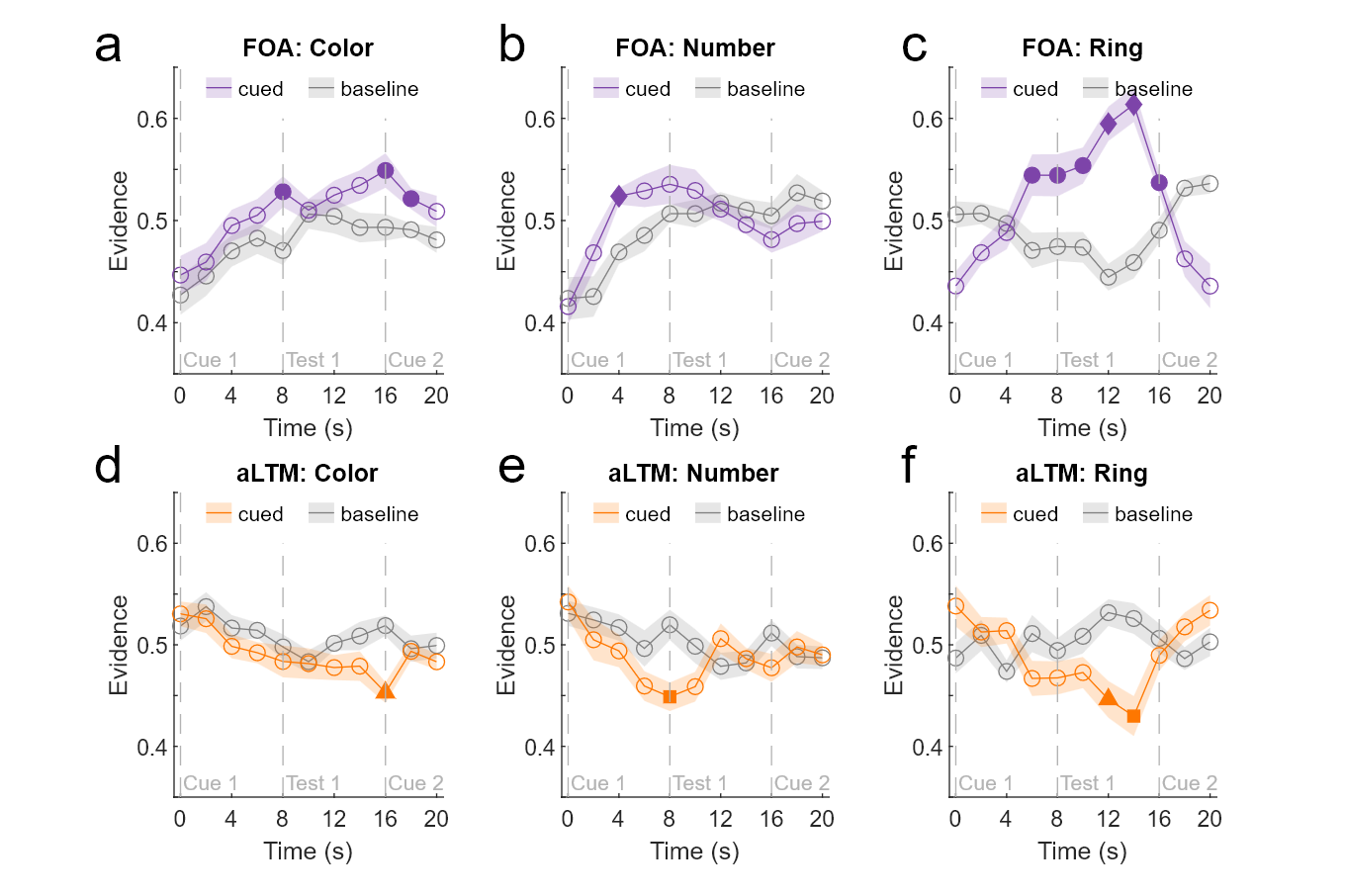

**Figure S5. Unthresholded importance maps for the FoA and aLTM classifiers**

Positive importance values (activity×weight) correspond to both the activity and weight being positive, while negative importance values correspond to both the activity and weight being negative. White voxels indicate regions of dropout or where the activity and weight were opposite signed (which was rare).

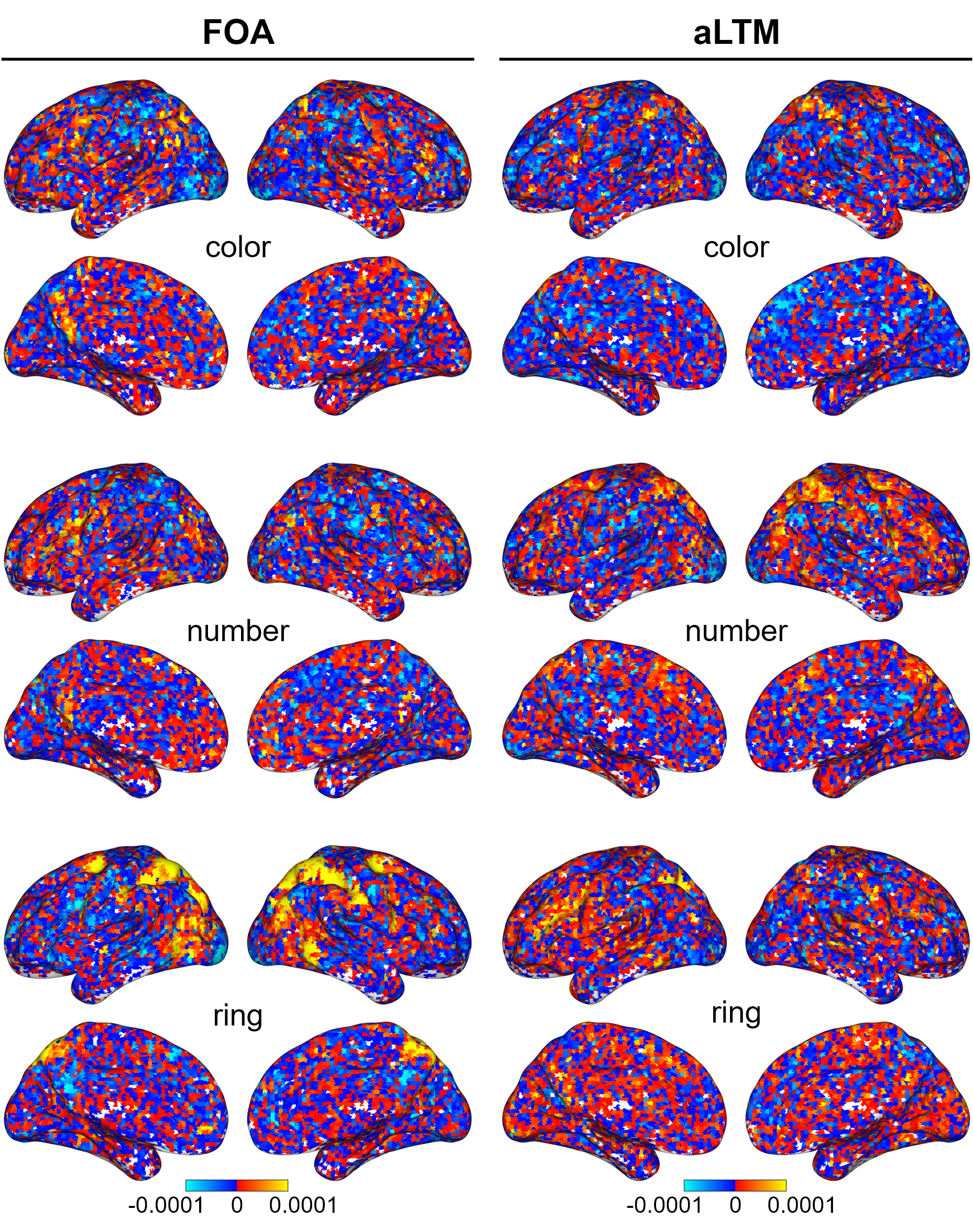

**Table S1. Significant classification results for individual time points**

| **Classifier** | **Figure panel** | **Time point (s)** | ***t*_15_** | ***p*** |
| --- | --- | --- | --- | --- |
| FoA accuracy for A1 | 2b | 4, 6, 8, 10 | 2.12, 4.99, 4.21, 4.18 | .026, <.001, <.001, <.001 |
| aLTM accuracy for A2 | 2e | 6, 8, 10 | 4.51, 2.97, 1.90 | <.001, .0048, .038 |
| FoA evidence for first-presented category across A1/A2 | 2c | 4, 6, 8, 10, 18, 20 | 2.96, 3.74, 3.49, 3.52, 2.08, 2.71 | .0048, .0010, .0016, .0015, .028, .0081 |
| FoA evidence for second-presented category across A1/A2 | 2c | 12, 14, 16, 18, 20 | 2.14, 4.15, 4.75, 4.50, 5.20 | .025, <.001, <.001, <.001, <.001 |
| aLTM evidence for first-presented category across A1/A2 | 2f | 2, 14, 16 | 2.09, 3.71, 1.76 | .027, .0011, .049 |
| aLTM evidence for second-presented category across A1/A2 | 2f | 4, 8 | 2.06, 2.31 | .029, .018 |
| FoA evidence for cued category during C1/T1/C2/T2 of repeat trials | 3a | 8, 10, 12, 14, 28, 30 | 2.77, 2.15, 2.88, 2.83, 2.76, 5.48 | .0071, .024, .0058, .0063, .0072, <.001 |
| FoA evidence for uncued category during C1/T1/C2/T2 of repeat trials^a^ | 3a | 0, 2, 4, 22, 24, 26 | 4.31, 4.55, 2.32, 2.41, 4.24, 3.26 | <.001, <.001, .035, .029, <.001, .0053 |
| FoA evidence (>baseline) for first cued category during C1/T1/C2/T2 of switch trials | 3b | 0, 2, 4, 6, 8, 10, 12, 14, 16 | 3.00, 2.59, 5.28, 4.11, 2.93, 1.93, 2.63, 1.90, 2.20 | .0045, .010, <.001, <.001, .0052, .037, .0095, .039, .022 |
| FoA evidence (vs. baseline) for first cued category during C1/T1/C2/T2 of switch trials^a^ | 3b | 20, 22, 24, 26, 28, 32 | 2.83, 2.25, 3.85, 3.88, 3.73, 4.47 | .013, .040, .0016, .0015, .0020, <.001 |
| FoA evidence (>baseline) for second cued category during C1/T1/C2/T2 of switch trials | 3b | 0, 2, 4, 22, 28, 30, 32 | 4.32, 3.46, 2.88, 2.82, 5.54, 2.20 | <.001, .0018, .0057, .0065, <.001, .022 |
| FoA evidence (vs. baseline) for second cued category during C1/T1/C2/T2 of switch trials^a^ | 3b | 8, 14, 18 | 2.71, 2.14, 2.82 | .016, .049, .013 |
| aLTM evidence for cued category during C1/T1/C2/T2 of repeat trials^a^ | 3c | 8, 14 | 2.94, 2.47 | .010, .026 |
| aLTM evidence for uncued category during C1/T1/C2/T2 of repeat trials^a^ | 3c | 0, 2, 36 | 3.07, 4.92, 2.50 | .0078, <.001, .025 |
| aLTM evidence for first cued category during C1/T1/C2/T2 of switch trials^a^ | 3d | 0, 14, 24 | 2.69, 2.54, 3.03 | .017, .023, .0085 |
| aLTM evidence for second cued category during C1/T1/C2/T2 of switch trials^a^ | 3d | 8, 10, 12, 14, 16 | 3.12, 2.26, 2.70, 2.43, 2.16 | .0071, .039, .017, .028, .047 |
| FoA evidence (>baseline) for cued category during C1/T1 | 4a | 4, 6, 8, 10, 12, 14, 16 | 3.47, 3.17, 3.22, 2.41, 3.99, 3.62, 1.95 | .0017, .0032, .0029, .015, <.001, .0013, .035 |
| aLTM evidence (<baseline) for cued category during C1/T1 | 4b | 6, 8, 10, 12, 14, 16 | 2.79, 2.77, 2.00, 2.37, 3.10, 2.81 | .0069, .0071, .032, .016, .0036, .0066 |
| FoA evidence (<baseline) for uncued category during C1/T1 | 4c | 6, 14, 18 | 3.97, 1.88, 1.77 | <.001, .040, .048 |
| aLTM evidence (>baseline) for uncued category during C1/T1 | 4d | 4, 6, 8, 10, 12, 16 | 2.20, 3.49, 1.94, 2.38, 2.10, 2.35 | .022, .0017, .036, .015, .027, .017 |
| FoA evidence (>baseline) for cued category during C2/T2 | 4e | 12, 14, 16 | 2.64, 7.44, 2.61 | .0093, <.001, .0099 |
| FoA evidence (<baseline) for uncued category during C2/T2 | 4e | 2, 4, 6, 8, 10, 12, 14, 16 | 1.96, 3.24, 3.88, 5.24, 5.72, 3.57, 1.78, 4.20 | .035, .0028, <.001, <.001, <.001, .0014, .048, <.001 |
| aLTM evidence (<baseline) for cued category during C2/T2 | 4f | 12 | 2.01 | .032 |
| aLTM evidence (> baseline) for uncued category during C2/T2 | 4f | 8 | 1.95 | .035 |

Time points are relative to the onset (0 s) of the stimulus of interest (A1, first array; A2, second array; C1, first cue; T1, first test; C2, second cue; T2, second test). For time courses spanning multiple stimuli of interest (e.g., A1 and A2), time points are labeled with respect to the onset of the first stimulus (i.e., A1 = 0 s, A2 = 8 s). Unless otherwise noted, all tests of classifier accuracy and classifier evidence corresponded to directional hypotheses and were based on one-tailed t-tests. ^a^These tests were two-tailed as they did not correspond to any a priori hypotheses.

**Table S2. Significant clusters from the searchlight classification analyses of FoA and aLTM**

| **Contrast and region label** | **# of voxels** | **Peak MNI coordinates (x, y, z)** | **Peak  *t*-value** |
| --- | --- | --- | --- |
| *FoA > chance* |  |  |  |
| L supramarginal gyrus (posterior) | 856 | -42, -43, 41 | 9.55 |
| L central opercular cortex | 104 | -45, 5, 11 | 8.05 |
| R parietal operculum cortex | 14 | 57, -28, 26 | 7.64 |
| R superior parietal lobule | 337 | 30, -52, 50 | 6.63 |
| L precentral gyrus | 62 | -24, -7, 50 | 6.60 |
| L middle temporal gyrus (temporo-occipital) | 21 | -54, -61, 11 | 5.80 |
| R frontal pole | 14 | 45, 38, 11 | 5.65 |
| R middle frontal gyrus | 19 | 30, -4, 56 | 5.52 |
| L lateral occipital cortex (inferior) | 10 | -45, -64, -4 | 5.35 |
| R middle temporal gyrus (temporo-occipital) | 13 | 45, -52, 5 | 5.17 |
| R precuneus cortex | 22 | 6, -61, 50 | 4.85 |
| *aLTM > chance* |  |  |  |
| L lateral occipital/superior parietal cortex | 62 | -21, -67, 44 | 6.59 |
| R lateral occipital/superior parietal cortex | 28 | 33, -61, 53 | 6.21 |
| L middle frontal gyrus | 10 | -45, 11, 32 | 5.27 |

Clusters were defined by a voxel-wise threshold of *p* < .001 (*t*_15_ > 3.733) and cluster extent of 10 voxels, corresponding to *p* < .05 (FWE-corrected). Labels are based on matching the peak coordinates with the Harvard-Oxford atlas.

**Table S3. Significant clusters from testing for overlap and differences across the searchlight classification analyses of FoA and aLTM**

| **Contrast and region label** | **# of voxels** | **Peak MNI coordinates (x, y, z)** | **Peak  *t*-value** |
| --- | --- | --- | --- |
| *FoA + aLTM (inclusively masked)* |  |  |  |
| L lateral occipital cortex (superior) | 10 | -30, -85, 26 | 8.03 |
| L lateral occipital/superior parietal cortex | 288 | -21, -73, 41 | 7.36 |
| L postcentral gyrus | 23 | -42, -37, 47 | 7.27 |
| L precentral gyrus | 22 | -45, 8, 29 | 6.63 |
| R superior parietal lobule | 138 | 30, -55, 53 | 6.17 |
| R middle temporal gyrus (temporo-occipital) | 10 | 54, -52, -4 | 3.80 |
| *FoA > aLTM* |  |  |  |
| R superior parietal lobule | 21 | 24, -49, 65 | 7.94 |
| L superior parietal lobule | 43 | -39, -40, 53 | 7.36 |
| R supramarginal gyrus (posterior) | 52 | 45, -37, 53 | 6.21 |
| R lateral occipital cortex (inferior) | 22 | 54, -64, 11 | 6.21 |
| L lateral occipital cortex (superior) | 13 | -24, -82, 23 | 6.21 |
| L superior parietal lobule | 12 | -30, -46, 65 | 5.37 |
| L lateral occipital cortex (superior) | 10 | -27, -82, 35 | 5.29 |
| L superior parietal lobule | 10 | -21, -58, 59 | 4.59 |
| *aLTM > FoA* |  |  |  |
| No significant clusters |  |  |  |

For the inclusive-masking procedure, the FoA and aLTM contrasts were individually thresholded at *p* < .01, resulting in a conjoint threshold of *p* < .001 (with an extent threshold of 10 voxels also applied). For the contrasts testing for differences between FoA and aLTM, clusters were defined by a voxel-wise threshold of *p* < .001 (*t*_15_ > 3.733) and an extent of 10 voxels, corresponding to *p* < .05 (FWE-corrected). Labels are based on matching the peak coordinates with the Harvard-Oxford atlas.
